# Aggregating multimodal cancer data across unaligned embedding spaces maintains tumor of origin signal

**DOI:** 10.1101/2025.05.14.653900

**Authors:** Raphael Kirchgaessner, Kaya Keutler, Layaa Sivakumar, Xubo Song, Kyle Ellrott

## Abstract

AI based embeddings offer the possibilities of encoding complex biological data into low dimensional spaces, called embedding spaces, that maintain the relationships between entities. There is an open question about the compatibility of embedding spaces that are created without any coordination. It has been assumed that signals in these unaligned embedding spaces would be destroyed if vectors were aggregated into summed values. We trained embedding models across different data modalities and tested aggregating the values together to test this assumption. Our research shows that signal from unaligned embedded values is conserved and able to still be used for learning tasks, such as data modality and tumor of origin recognition.

## Motivation

As AI techniques are applied to biomedical and genomic data, one of the first questions is how the data is encoded and represented, as the first step in many AI based pipelines is encoding the source data into an embedding space [1], [2], [3] . Embedding spaces are lower dimensional manifolds where relative relationships of elements are maintained, for example similar elements are placed closer to each other. In modern machine learning, combining diverse data types into single embedding vectors, using aggregation functions such as summation, is a common practice [4], [5]. These aggregations significantly enhance model performance in areas such as natural language processing, genomics and computer vision [6], [7], [8], [9]. This fusion of heterogeneous data allows for richer representations, capturing multiple facets of complex datasets. However, this process often obscures the origins of the data, making it challenging to interpret or analyze the specific contributions of individual modalities [10]. Understanding the sources of these combined embeddings is crucial for improving model interpretability and ensuring that critical information is preserved.

Importantly, there is an open question about how the aggregation of non-aligned embedding vectors will affect recognition. In creating embedding spaces different dimensions tend to encode complex concepts, with the relative value associated with the position in the axis. In language models, concepts such as gender or the concept of royalty can be associated with different axes. The difference between the words ‘king’ and a ‘queen’ moves along the gender axis, while the difference between ‘man’ and ‘king’ moves along the royalty axis. Importantly, because these different axes were created in coordinated optimization, the summation of two vectors, each one representing movements in singular axis, produce a coherent combined effect, ie adding a vector in the direction of gender to one with the direction of royalty will move from the word ‘man’ to ‘queen’ [11]. This property has widely been taken advantage of in LLM and graph neural networks. In the case of multi-modal data, where embedding spaces have been created with no consideration of other data encoding, there is no guarantee that the summation of vectors will not cause interference between concepts encoded in different spaces. There is the possibility that if the major axes in different embedding spaces conflict with each other, any recognizable signal could be lost in the aggregation. We proposed a set of experiments to measure the effect of interference between different embedding spaces when aggregated together.

Biomedical data related to precision oncology comes in a variety of different forms including genomic and transcriptomic profiles, imaging data and clinical notes [12], [13], [14]. One strategy would be to create a multi-modal embedding space, where all data modalities are commonly encoded, using a method such as the Contrastive Language-Image Pre-Training (CLIP) algorithm [15] which helped to build the text to image conversion of DALL-E [16]. However, we were interested to see if a more simplistic strategy for multi-modal embedding integration could be deployed for enabling search, clustering and recognition of connected cancer records. Our primary motivation was to investigate whether it is possible to combine heterogeneous patient data—such as RNA transcription levels, H&E images, and patient clinical text annotations—into a unified embedding while still retaining the ability to distinguish the sources that contributed to the aggregated vector. Importantly, we wished to determine if this analysis could be done without coordinating the various embedding manifolds, meaning that each of the various embedding spaces are created independently and only organized using the uni-modal source information.

Graph learning algorithms have been proposed as effective tools for encoding graph-structured information [17], [18]. However, most existing methods have been demonstrated using graphs constructed from a single data modality. In the case of complex biological data, there may be multiple data modalities, and thus multiple embedding spaces, that can be connected together in a structured graph. In order for these methods to be effective, it must be possible to aggregate vectors from multiple data modalities, without losing all relevant information to noise. This is possible in single-modality embedding spaces, however whether or not this capacity is maintained when aggregating across uncoordinated embeddings is less well known.

In addition, we evaluated whether the summed embeddings maintain enough information integrity to be stored and efficiently retrieved from vector databases. Use of embedding methods for creating vector indices of data allows for semantically aware search of document stores. Metrics such as cosine similarity, Euclidean distance, and dot product were used to assess whether the aggregated embeddings could still be meaningfully compared and queried. This aspect of the study is highly relevant, as the ability to store and retrieve embeddings in vector databases is critical for real-world applications, including personalized medicine, where fast, reliable retrieval of patient-specific data can lead to better treatment outcomes.

Lastly, we aimed to assess whether these summed embeddings, generated from heterogeneous multi-modal data, could effectively classify cancer types. This would demonstrate that despite the complexity of the data fusion, the resulting embeddings still contain enough discriminative power to distinguish between different cancer types. This investigation could have significant implications for cancer diagnostics, enabling more accurate and interpretable classification models in the future.

## Approach

To investigate whether simple aggregation of embeddings from unaligned latent spaces can yield meaningful representations for downstream tasks, we developed two complementary sampling and evaluation strategies.

The first strategy focused on modality-level sampling to address the composition recognition problem. In this setting, the goal was to determine whether it is possible to infer which data modalities (e.g., RNA, H&E, clinical annotations) were used to construct a given aggregate vector. To generate the training data for this task, we randomly sampled embeddings from different modalities without regard to patient identity, and combined them to form synthetic aggregate vectors. A recognition model was then trained to predict the constituent modalities present in each input vector. This approach allowed us to assess whether naive aggregation preserves sufficient modality-specific structure to support decomposition, despite the embeddings originating from unaligned latent spaces.

The second strategy was designed to assess the clinical and biological utility of the aggregated embeddings. Here, the task was to predict cancer-relevant attributes—including cancer type (e.g., BRCA, BLCA), tumor mutational burden (TMB), and molecular subtype—based on patient-level aggregate vectors. To construct these vectors, we performed patient-level sampling, where each aggregate vector was composed of heterogeneous embeddings corresponding to the same individual. These included representations derived from RNA-seq profiles, H&E images, clinical annotations, and somatic mutation data. This setup enabled us to evaluate whether the aggregated vectors retained enough biological signal to be useful for clinically relevant prediction tasks.

All embeddings were generated using either Variational Autoencoders (VAEs) or Sentence-BERT (sBERT) (Figure 1a). Each embedding represented a datapoint tied to a specific patient or modality. The only coordination between different embedding models was dimensionality alignment: all vectors were projected into a shared 768-dimensional space to match the output size of the sBERT model.

**Figure 1:**
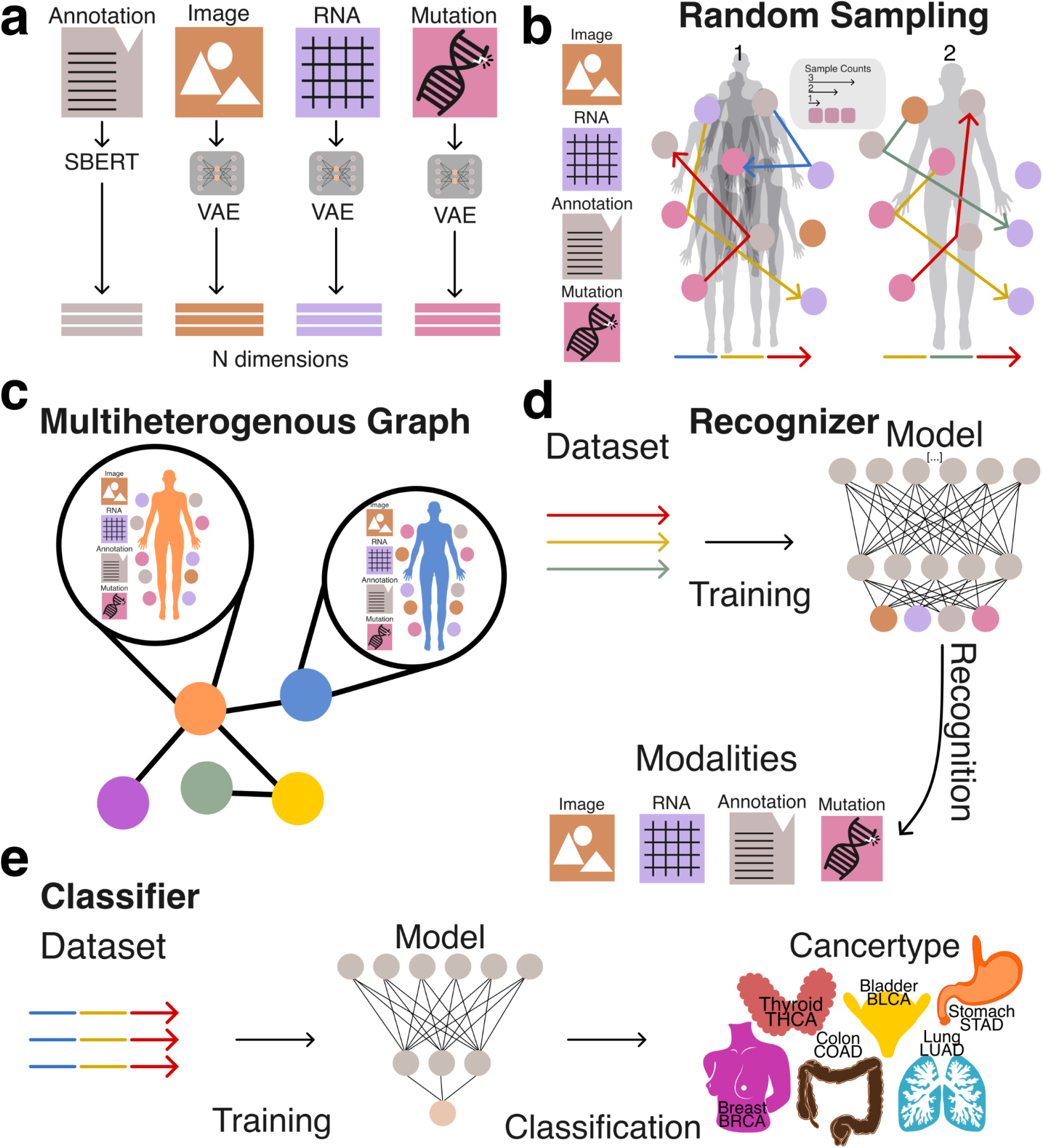
Overview of embedding generation and network architecture. **A:** Modalities (RNA, Image, Mutation & Annotation) used to generate embeddings by using either SBERT or a VAE. **b:** Illustration of random samplings performed over multiple patients (1) and one patient (2), demonstrating different sampling counts. **c:** Schematic representation of a heterogeneous graph incorporating patients and their associated modalities. **d:** Schematic of the recognizer experimental setup to recognize multiple modalities, including image, RNA, annotations, and mutation data. **e:** Schematic of the classifier experiment setup which predicts the patient’s cancer type, classified as BRCA, LUAD, BLCA, THCA, COAD, or STAD.

To prepare the embeddings for use with traditional machine learning pipelines and vector databases, we applied structured random sampling and aggregation procedures. This transformation was critical for enabling efficient storage, retrieval, and analysis of heterogeneous data types using conventional vector-based methods. By encapsulating diverse multimodal representations into a unified vector format (Figure 1b, 1c), we facilitated systematic evaluation of whether naive aggregation preserves informative structure across both tasks.

Together, these two strategies allowed us to evaluate the extent to which simple vector aggregation from unaligned modality-specific embeddings can support downstream machine learning applications, including modality composition recognition and clinically meaningful phenotype prediction (Figure 1d, 1e).

## Results

### Unsupervised clustering

To evaluate the intrinsic separability of cancer types without relying on complex or computationally intensive deep learning models, we applied a Gaussian Mixture Model (GMM) clustering algorithm to patient-level aggregate embeddings generated via the second sampling strategy (patient-level sampling) (Figure 2a). GMM is an unsupervised learning method that models data as a mixture of Gaussian distributions, allowing it to group similar data points by estimating their underlying probabilistic structure. The number of GMM components was set equal to the number of cancer types in the dataset (BRCA, BLCA, STAD, COAD, THCA, LUAD), enabling a direct assessment of whether embeddings corresponding to different cancer types could be naturally grouped.

**Figure 2:**
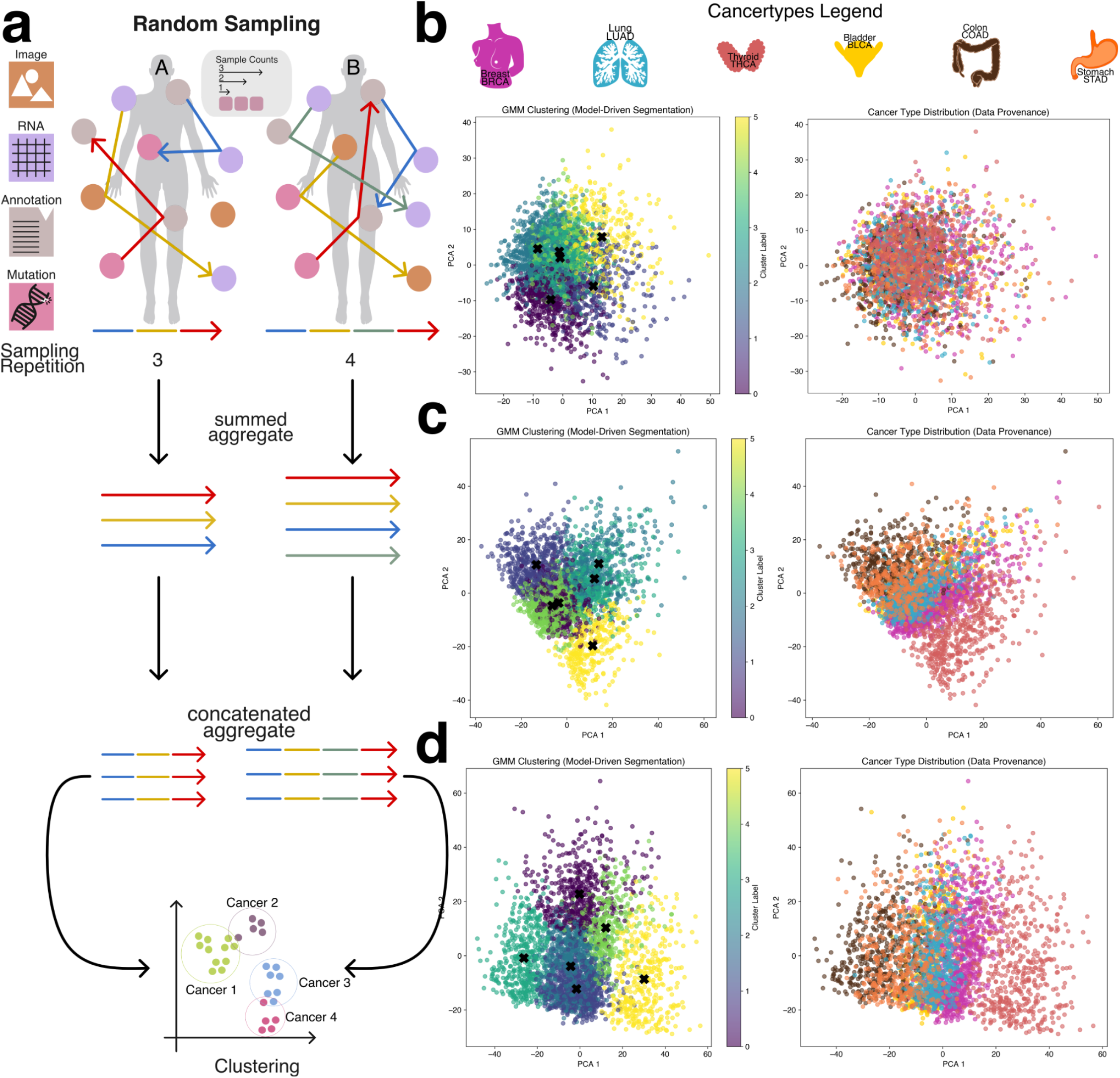
Unsupervised clustering of summed and concatenated embeddings. **a:** Schematic depiction of patient-specific embeddings obtained by aggregating and concatenating vectors derived from random samplings. Random samplings performed on the graph structure generate vectors concatenated into a single embedding vector for subsequent usage in a gaussian mixture model (GMM) clustering. **b:** PCA representation of aggregated and contacted embeddings using a sample count of three and a sample count repeat of three, shows little differentiation between cancer types. **c:** Increasing both sample count and repetition by one to four both PCA and GMM clustering shows a clearer separation between cancer types. **d:** Using a sample count and repeat of 5 further enhances cancer type separation with THCA (far right cluster) observing the strongest separation.

Using a low sampling configuration—specifically, a sample count of three and sampling repeats of three—the resulting clusters were largely indistinct, with significant overlap between cancer types and no clear boundaries between groups (Figure 2b). However, as both the sample count and sampling repeats increased, clustering quality improved markedly. Increasing the number of modality embeddings used in aggregation (i.e., higher sample counts) and repeating the sampling process multiple times allowed the aggregated embeddings to better capture patient-specific variation and inter-modality relationships. This led to progressively clearer separation between cancer types (Figures 2c and 2d), with THCA emerging as the most distinctly clustered cancer type, while other cancers such as BLCA, STAD, and COAD exhibited closer proximity and partial overlap.

The clearest clustering performance was achieved with a sample count of five and five sampling repeats, which resulted in well-defined cluster boundaries and improved alignment between GMM components and cancer labels (Figures 2b–d, Supplemental Figures 1–3). Notably, when cancer types appeared between two GMM clusters, they often exhibited biological similarities to both, reinforcing the non-linearity of the underlying molecular patterns. These results demonstrate that even in the absence of supervised learning, cancer type identity is at least partially encoded in the aggregated embedding space, and that unsupervised clustering performance is strongly influenced by the richness and redundancy introduced through the sampling process.

### Identification of Component Embeddings in Aggregated Representations

In this portion of the study, we evaluated the ability of machine learning models to accurately identify the underlying data modalities and cancer types used to generate each aggregate embedding. Successful recognition of the constituent modalities and cancer type supports our hypothesis that, despite the naive combination of embeddings, sufficient structural information is preserved to enable meaningful downstream analyses. To support this hypothesis we implemented two modeling strategies: a simple approach, where the model was tasked with distinguishing between the base modalities (e.g., RNA, image, mutations, and annotations), and a cancer-specific approach, which further required the model to identify the cancer type from which the embeddings were derived (Figure 3a).

**Figure 3:**
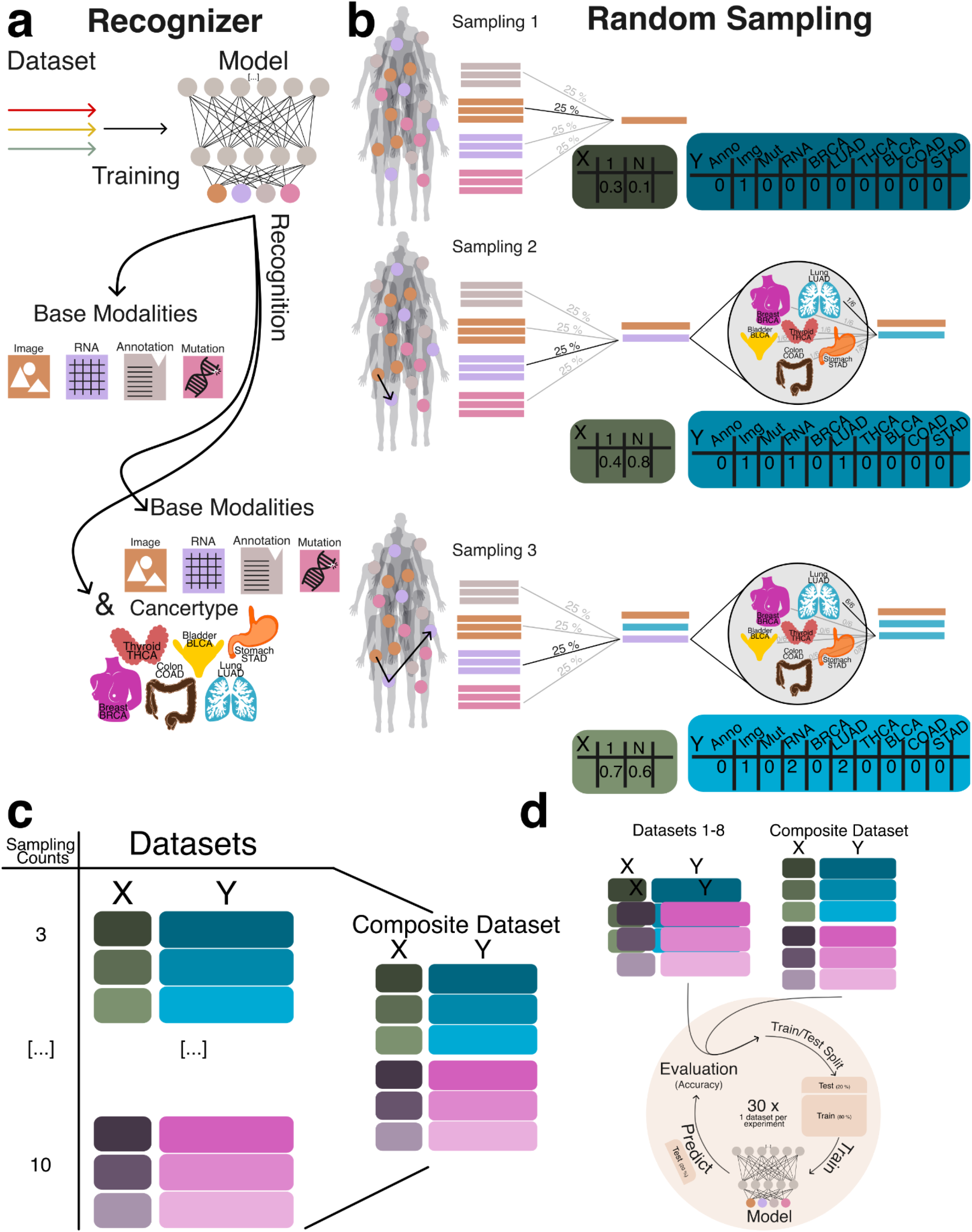
Embedding generation for the recognizer network. **a:** Recognizer setup including a simple recognizer trained to identify base modalities (RNA, image, somatic mutations, and annotations), and a cancer-specific recognizer designed to additionally distinguish the cancer type from which embeddings are derived. **b:** Random Sampling (RS) process shown for a sample count of 3. The process begins by selecting a random embedding (X) and recording its modality and cancer type in Y. A second embedding is sampled from the same cancer type, summed with the first, and tracked. A third embedding is then added, completing the aggregated vector while continuing to track contributing modalities. **c:** Composite dataset generation by combining all datasets generated using sample counts ranging from 3 to 10. **d:** Both isolation and composite datasets are used for training and evaluating model performance, with training and testing conducted using at least n > 30 independent runs per condition.

To generate datasets for training and evaluating the recognition model, we created datasets ranging from three to ten random sampling steps, each containing 15,000 data points (Figure 3b). To assess model generalizability, we constructed a composite dataset by integrating all datasets containing between three and ten constituents, resulting in a total of 90,000 summed embeddings (Figure 3c). To establish a baseline, we employed a multiclass logistic regression model (MCRM) to evaluate the feasibility of recognizing the composition of aggregate embeddings, aiming to determine whether the model could accurately distinguish the base modalities used to generate each aggregate vector and to evaluate whether more complex models are necessary. Additionally, we developed a deep learning model (DL) to leverage a more expansive feature space for representation learning. Both models were trained and evaluated on datasets split into training and test sets, with each experiment count of n >= 30 to ensure robustness (Figure 3d). Given the dataset’s class imbalance and zero inflation, we evaluated model performance using the Matthews correlation coefficient (MCC), balanced accuracy, and F1 score. MCC was chosen as the primary metric due to its suitability for imbalanced data [19], [20]. Figure 4a illustrates model performance under the simple aggregation approach, which corresponds to the first strategy described earlier. In this setting, we trained isolated models, where each model was exposed only to embeddings generated using a fixed number of randomly sampled modality vectors (referred to as sample count, e.g., three). Sample count serves as a proxy for aggregation complexity, with higher counts introducing more heterogeneity into each embedding. Both the MCRM baseline (BL) and deep learning (DL) models achieved high Matthews correlation coefficient (MCC), accuracy, and F1 scores at lower sample counts. However, as the number of aggregated vectors increased, performance differences became more pronounced—particularly for somatic mutation and annotation embeddings. The MCC for the MCRM BL model declined below 0.8 for somatic mutations and below 0.6 for annotations, while the DL model maintained stable performance close to 0.9 across all sample counts.

**Figure 4:**
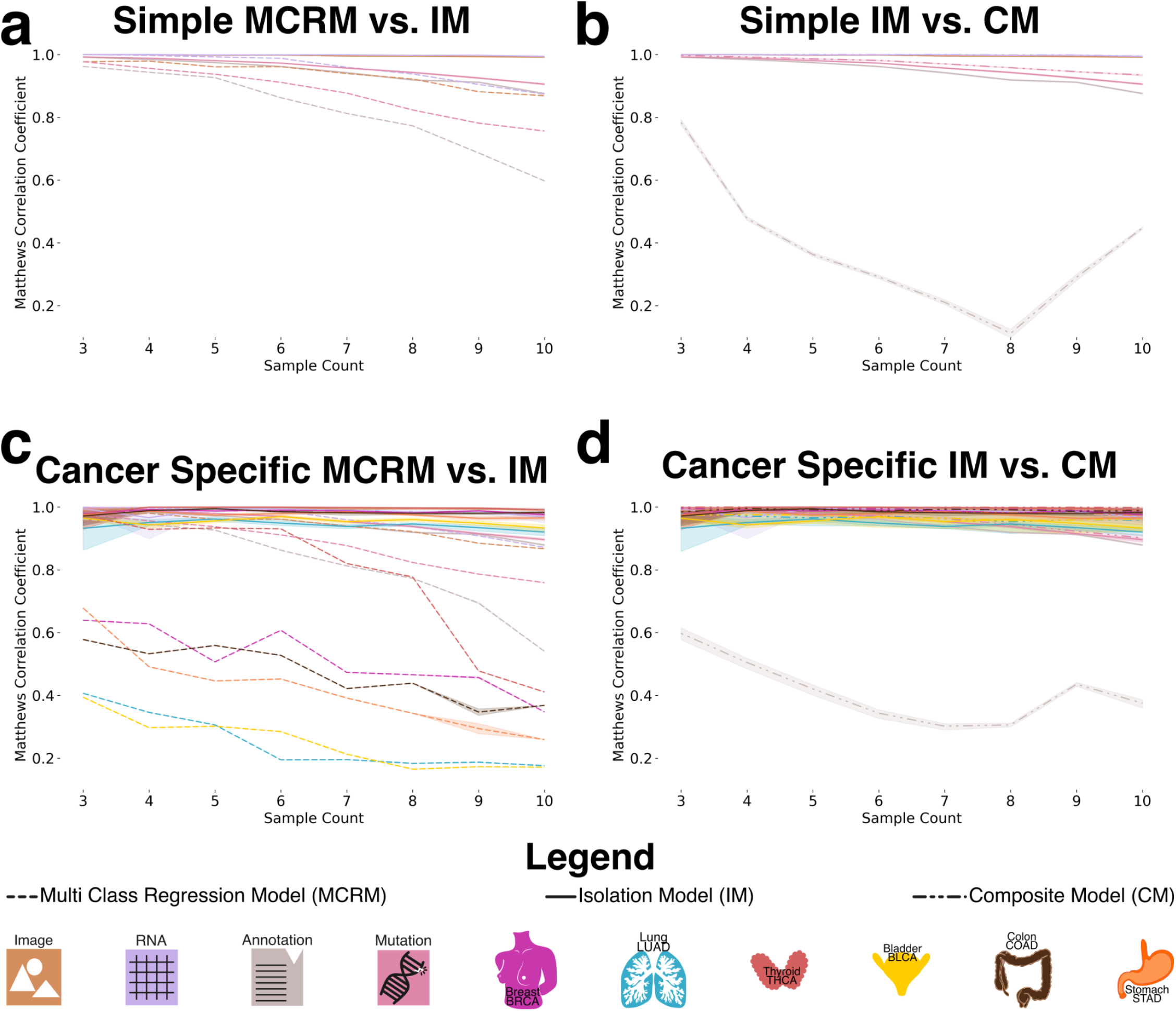
Performance of deep learning and baseline models across aggregation strategies and sampling contexts. **a:** Performance of the simple aggregation approach using isolation models (IM) across modalities, evaluated at sample counts ranging from 3 to 10. The multi-class logistic regression model (MCRM), used as a baseline, achieves MCC values between 0.6 and 0.8, while the DL-based IM consistently outperforms it, achieving MCC values above 0.9. **b:** Comparison between composite models (CM) and isolation models (IM) under the simple aggregation setting. Performance remains high across most modalities for both models, except for the annotation modality, where the CM shows a marked decline in MCC with increasing sample count, dropping as low as 0.4. **c:** Performance under the specific-cancer approach comparing cancer-specific IMs to the MCRM baseline. The baseline shows poor performance for several cancer types, particularly BLCA, LUAD, and STAD, with MCC values between 0.2 and 0.4. In contrast, the DL-based IMs achieve strong and consistent performance across all cancer types (MCC > 0.9). **d:** Comparison between cancer-specific composite models (CM) and isolation models (IM). While most modalities maintain high performance (MCC > 0.85) across both models, the annotation modality shows poor performance in the CM across all sample counts, consistent with trends observed in panel b.

Figure 4b compares two deep learning models: an isolation model, trained separately on embeddings generated from a fixed sample count, and a composite model, trained on a dataset comprising all sample counts. Across most modalities, both models performed similarly. However, for annotation embeddings, the composite model consistently underperformed relative to the isolation model. Specifically, the composite model’s MCC ranged between 0.2 and 0.6, while the isolation model maintained higher and more stable MCC values, demonstrating greater robustness when trained on uniformly structured inputs. We hypothesize that this discrepancy stems from both the nature of the embeddings and the interaction between embedding heterogeneity and aggregation complexity. Notably, the annotation embeddings were generated using Sentence-BERT (sBERT), whereas all other modality embeddings were generated using Variational Autoencoders (VAEs). As a result, the annotation embeddings originate from a fundamentally different and more semantically structured latent space, making them less compatible with VAE-based embeddings during aggregation. This heterogeneity likely contributes to the model’s difficulty in learning consistent representations when multiple annotation embeddings are combined in a composite setting. At lower sample counts (e.g., three), the aggregation of a small number of heterogeneous embeddings may still preserve relatively coherent inter-modality patterns, enabling the model to extract useful signals despite differences in latent space. However, as the sample count increases, feature incompatibility across modalities becomes more pronounced. The model struggles to reconcile divergent feature distributions and representations not designed to be directly comparable, leading to a marked decline in performance as the sample count increases from 3 to 8. Interestingly, performance begins to recover at higher sample counts (around nine or more), suggesting that sufficient aggregation of diverse embeddings may allow the model to detect higher-order patterns across modalities. At this point, redundancy and complementary information across heterogeneous embeddings may help the model abstract away modality-specific noise and form more robust decision boundaries. Certain cancer-related features may become detectable across multiple modalities, allowing the model to generalize effectively despite latent space mismatches.

Figure 4c presents results from the specific-cancer approach, where isolated models were trained using a fixed number of samples per cancer type (e.g., three samples per cancer). Despite this constraint, the DL model maintained stable performance across cancer types. In contrast, the MCRM BL model exhibited significant performance degradation under these conditions, with the effect particularly pronounced for cancer types such as lung adenocarcinoma (LUAD) and bladder cancer (BLCA), where MCC dropped to 0.2 when trained on a sample count of 10.

Figure 4d mirrors the composite versus isolation comparison within the specific-cancer context, again evaluating only deep learning models. Consistent with the findings in Figure 4b, annotation embeddings exhibited the most pronounced performance drop in the composite setting. The composite model failed to achieve MCC values above 0.6 for annotation embeddings, with performance falling below 0.4 for sample counts between 5 and 8. In contrast, the isolation model maintained consistently high performance, with MCC values remaining above 0.9 across all sample counts.

We attribute this pattern to the same factors observed in Figure 4b, with an important clarification regarding the nature of the underlying embedding spaces. Annotation embeddings are generated using sBERT, which produces deterministic embeddings—each input is mapped to a fixed point in a high-dimensional semantic space without an explicit distribution. In contrast, embeddings from other modalities are derived from Variational Autoencoders (VAEs), which learn probabilistic latent spaces and encourage structured, continuous representations. The VAE architecture supports smooth interpolation and aggregation of latent vectors, properties that are not inherently preserved in the sBERT embedding space.

As a result, when annotation embeddings are combined with VAE-based embeddings in the composite setting—especially across varying sample counts and cancer types—the lack of compatibility between these embedding spaces likely introduces feature conflicts that degrade model performance. Isolation models, by contrast, operate on more uniformly structured inputs and are thus less affected by this latent space mismatch, enabling them to maintain stable performance even when incorporating heterogeneous modality types.

To further evaluate model robustness, we introduced a controlled noise perturbation framework in which a defined proportion of modality embeddings within each aggregated vector was systematically replaced with random Gaussian noise. This procedure preserved the total number of embeddings per vector—maintaining the original sample count—and enabled assessment of the model’s ability to differentiate true modality embeddings from noise. The deep learning (DL) model was trained exclusively on clean (noise-free) data and subsequently evaluated on perturbed datasets. Performance was assessed using Matthews correlation coefficient (MCC), F1 score, and balanced accuracy to quantify the impact of noise on the model’s ability to correctly infer embedding composition (Figure 5a).

**Figure 5:**
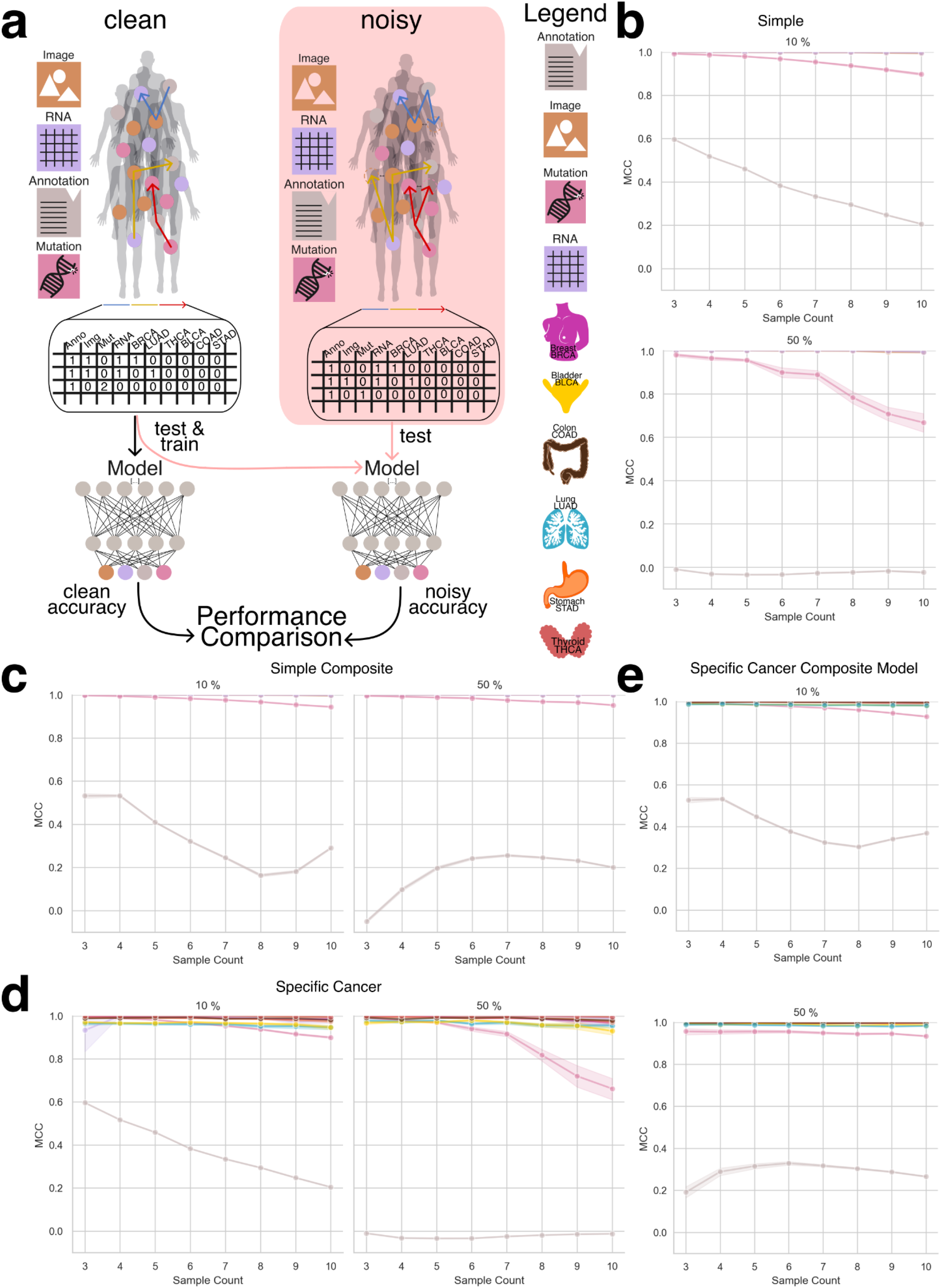
Performance of deep learning recognizer models under controlled noise perturbation. **a:** Overview of the experimental setup: models were trained on clean (noise-free) aggregated embeddings and evaluated on test data with controlled Gaussian noise introduced at 10% and 50% of embedding positions. Noise was applied while preserving the original sample count to assess robustness in distinguishing true modality embeddings from noise. **b:** Performance of the simple aggregation approach using isolation models (IM) under noisy conditions. At 10% noise, the model maintains high MCC values for RNA and image modalities, while performance for the mutation modality decreases moderately and the annotation modality shows poor performance even at low noise levels. At 50% noise, annotation performance remains consistently low, and mutation modality performance declines progressively with increasing sample count. **c:** Performance of the specific-cancer approach using isolation models (IM) at 10% and 50% noise. Models demonstrate stable MCC values across most modalities at 10% noise, with the exception of annotations. At 50% noise, annotation performance further deteriorates, and mutation embeddings exhibit notable decline for sample counts greater than six. **d:** Performance of the simple aggregation composite model (CM) under noisy conditions. Despite the presence of noise, the model achieves high MCC values (>0.9) for most modalities at both 10% and 50% noise. The annotation modality continues to underperform across all conditions. Notably, mutation embeddings show improved stability compared to the simple isolation model. **e:** Performance of the specific-cancer composite model (CM) under 10% and 50% noise. The model maintains strong MCC performance across most modalities and cancer types, except for the annotation modality, which consistently exhibits poor performance across both noise levels.

Due to the MCRM baseline model’s limited performance in previous experiments, this analysis was conducted exclusively with the DL model. We evaluated both the simple aggregation approach and the specific-cancer approach, comparing model performance across increasing noise levels. Figures 5b–e present results across noise conditions ranging from 10% to 50% embedding replacement.

Under the simple aggregation approach using isolation models (Figures 5b and 5c), performance remained robust at lower noise levels. Mutation embeddings, in particular, demonstrated high resilience, maintaining MCC values above 0.8 when 10% of the input was replaced with noise. However, at 50% noise, mutation performance declined to approximately 0.6 MCC, suggesting reduced but still meaningful predictive capacity. In contrast, annotation embeddings—derived from the sBERT model rather than VAE-based encoders—exhibited consistently lower MCC values, even under minimal noise, reflecting their greater sensitivity to perturbation and underlying latent space incompatibility.

In the specific-cancer setting using composite models (Figures 5d and 5e), annotation embeddings continued to perform poorly across all noise levels. However, mutation embeddings showed enhanced stability compared to the isolation setting, with MCC values exceeding 0.9 even at higher noise levels. This suggests that composite training across a broader dataset may help the model generalize more effectively in the presence of noise, at least for certain modalities.

These results underscore the DL model’s resilience to input perturbations and highlight the importance of both aggregation strategy and embedding origin in determining robustness. While noise degrades performance in a modality-dependent manner, the ability to retain high MCC values—particularly in the mutation modality—demonstrates the potential of deep learning to support reliable composition recognition even under challenging conditions.

### Tumor of Origin Identification

The classification task in this study aimed to predict specific cancer types from aggregated embeddings. While the dataset remained consistent with previous experiments, this task employed the second sampling strategy—patient-level sampling—to ensure that each aggregated embedding corresponded uniquely to a single patient. Each embedding integrated heterogeneous modality representations, including RNA-seq, H&E image features, clinical annotations, and somatic mutation profiles, reflecting the structure of real-world clinical datasets where multi-modal information is available and is ideally combined for tasks such as cancer type prediction.

To construct the aggregated embeddings, we performed multiple rounds of random sampling (sample repetitions) per patient. In each round, a subset of that patient’s modality-specific embeddings was randomly selected and summed to produce a single embedding of fixed dimensionality (e.g., 768 dimensions). To capture additional intra-patient variability and enhance representational richness, this sampling process was repeated multiple times. The resulting sampled embeddings were then concatenated to form a final, patient-specific aggregate embedding. For example, if three sampling rounds were performed, the final embedding would have a dimensionality of 3 × 768 = 2304. This approach preserved both the diversity of intra-patient modality contributions and a standardized input structure suitable for downstream machine learning analysis (Figure 6a).

**Figure 6:**
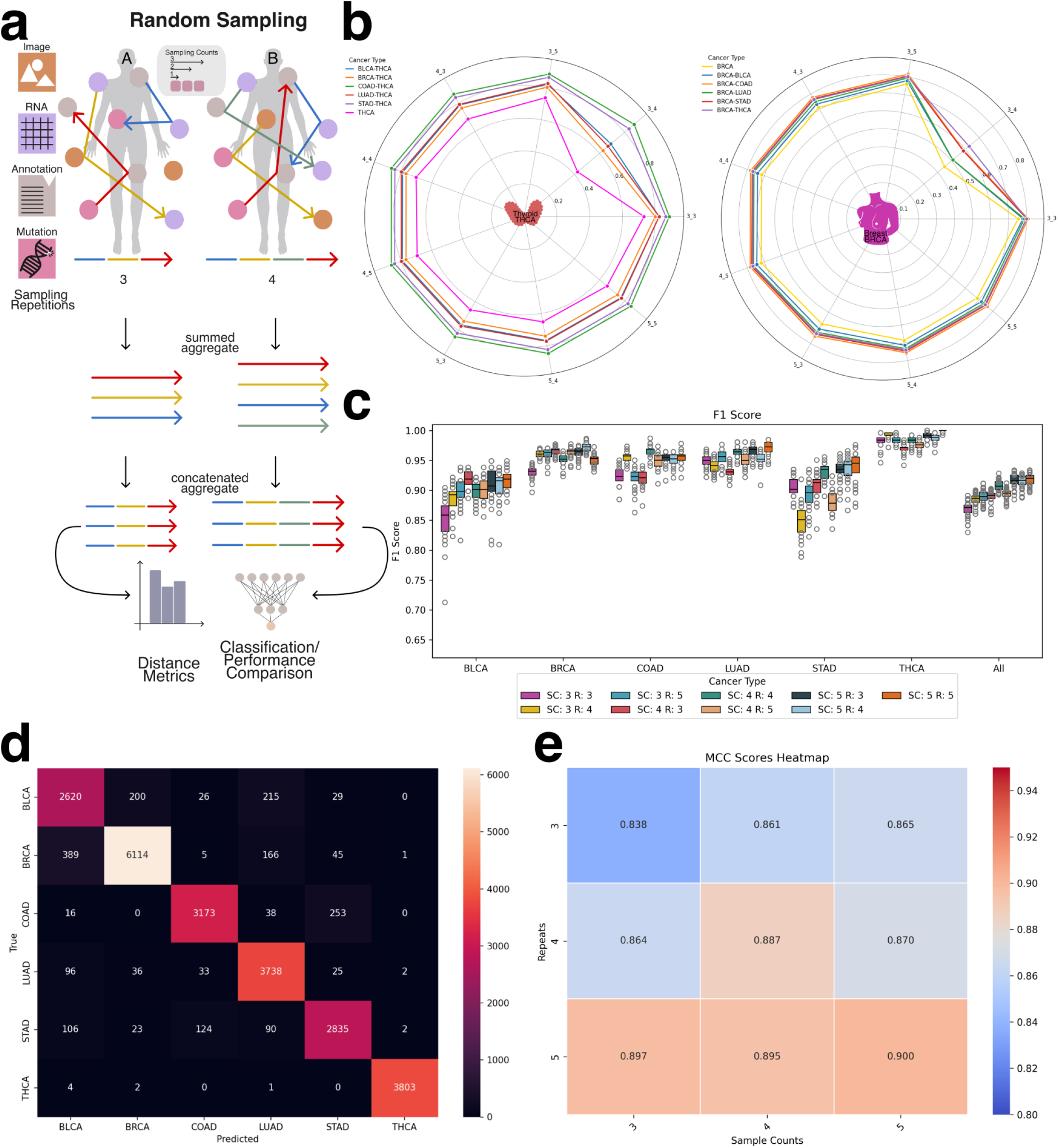
Classifier configuration and traditional distance metrics. **a:** Schematic depiction of patient-specific embeddings obtained by aggregating and concatenating vectors derived from random samplings. Random samplings performed on the graph structure generate vectors concatenated into a single embedding vector for subsequent distance metric calculations. **b:** Intra- and inter-cluster distances for THCA, demonstrating smaller intra-cluster distances relative to inter-cluster distances. Intra- and inter-cluster distances for BRCA, similarly illustrating smaller intra-cluster distances compared to inter-cluster distances. **c:** Boxenplots depicting F1 score distributions across multiple cancer types for varying combinations of sample counts (SC) and repetitions of sample counts (R). Classification performance was consistently high (>0.75), with slightly lower values observed for BLCA and STAD. **d:** Confusion Matrix depicting true positive and false positive predictions for all classes of the classifier. **e:** Heatmap representation of MCC scores demonstrating increased classifier performance with greater walk distances and walk amounts.

Before evaluating classification models, we conducted an exploratory analysis using traditional vector similarity metrics—including Euclidean distance, cosine similarity, and dot product—to assess whether the aggregated embeddings inherently reflect cancer-type-specific structure. This analysis was motivated by the widespread use of such metrics in vector database retrieval systems, where embedding similarity governs search and retrieval operations. By comparing intra-cancer and inter-cancer distances, we evaluated whether these metrics alone could meaningfully differentiate between cancer types. As shown in Figure 6b and Supplemental Figure 4, patient embeddings from the same cancer type exhibited higher similarity to each other than to embeddings from other cancer types. These findings indicate that even in the absence of supervised learning, aggregated embeddings retain informative structure relevant to cancer identity and may be applicable in retrieval-based clinical systems.

### Tumor classification using a deep learning model

For deep learning-based classification, the concatenated aggregate embeddings served as input features to the classifier network. These embeddings integrated information obtained from multiple random samplings of each patient’s modality-specific data, capturing both the heterogeneity and redundancy across input modalities. Model performance, evaluated using F1 score (Figure 6c), demonstrated the effectiveness of this approach in predicting cancer types from multi-modal patient-level embeddings. Across all cancer types, the model consistently achieved F1 scores ≥ 0.8 and accuracy scores ≥ 0.7. The lowest performance was observed for bladder cancer (BLCA) and stomach adenocarcinoma (STAD). As shown in the confusion matrix (Figure 6d), misclassifications frequently occurred between LUAD and BLCA, as well as between COAD and STAD, consistent with previously reported molecular overlaps (e.g., https://pmc.ncbi.nlm.nih.gov/articles/PMC5965261/).

To investigate the influence of sampling design, we evaluated how model performance varied with different sample counts and sampling repetitions. As shown in Figure 6e (MCC) and Supplemental Figure 5 (F1 score), increasing either the number of embeddings per sample or the number of sampling rounds consistently improved classification performance. These results suggest that aggregating additional heterogeneous information from a patient—regardless of the specific modality—enhances patient-level representation and improves the model’s ability to differentiate between cancer types.

To extend this framework to additional predictive tasks, we evaluated the feasibility of using the same aggregated embeddings to classify cancer subtypes (Supplemental Figures 6 and 7) and to predict tumor mutational burden (TMB) (Supplemental Figures 8 and 9). The model achieved reasonable performance across both tasks, further supporting the utility of integrating multiple unaligned modalities to construct a robust, patient-specific embedding suitable for a wide range of clinically relevant predictions.

## Discussion & Future Work

Our findings demonstrate that the performance of both the recognizer and the classifier model indicates that embeddings from diverse embedding spaces can be effectively combined to describe local subgraphs without the need of a shared latent space. This approach enables the successful identification of both the composition of aggregated embeddings and the cancer type, sub type or tumor mutational burden associated with them.

Furthermore, our findings demonstrate that in a heterogeneous graph, aggregates covering specific regions can serve as a secondary index, enabling the linkage of multimodal records to individual entities. Furthermore, our results highlight the feasibility of using heterogeneous networks for graph neural network transformations, expanding the potential applications of these architectures. Vector databases have gained significant attention as the foundation of Retrieval-Augmented Generation (RAG). Algorithms such as Hierarchical Navigable Small Worlds (HNSW) [PMID:30602420] allow for the rapid indexing of vector based data. This work demonstrates that despite mixing uncoordinated embedding spaces, aggregate vectors still contain sufficient data to be identifiable regarding their modality composition. This will allow for indexing of complex heterogeneous data graphs, not only at the per node level, but also at the neighborhood level. Records in a complex patient information system can be rapidly compared, even if the assays available are not equivalent between each individual.

We hypothesize that the observed performance degradation in the annotation modality stems from fundamental differences in the embedding generation methods. While all other modalities were encoded using variational autoencoders (VAEs), annotation embeddings were generated using Sentence-BERT (sBERT). This architectural mismatch likely contributed to the reduced performance of the annotation embeddings, particularly in recognition tasks and under composite training conditions. VAEs are designed to construct a continuous, structured latent space by enforcing a probabilistic distribution over the learned representations. This structure promotes smooth interpolation, supports meaningful aggregation, and enhances compatibility across embeddings—properties that are critical for robust downstream learning. In contrast, sBERT produces deterministic embeddings optimized for semantic similarity, without enforcing continuity or geometric smoothness in the latent space. As a result, sBERT-derived embeddings may exhibit a more fragmented or irregular structure, with weaker alignment to the representations learned by VAEs. This mismatch likely becomes more problematic as the sample count increases in aggregated embeddings, introducing feature conflicts that the model struggles to reconcile. Specifically, when a small number of embeddings (e.g., sample count of three) are combined, the model may still learn relatively coherent patterns between modalities, even in the presence of latent space heterogeneity. However, as additional embeddings are aggregated, the lack of alignment between the sBERT and VAE embedding spaces amplifies inconsistency, degrading performance—particularly in composite models where embeddings from a wide range of sample counts and cancer types are mixed. Interestingly, at higher sample counts (e.g., nine or more), we observed a partial recovery in model performance. We hypothesize that this is due to the emergence of higher-order patterns and redundancies across modalities. As more diverse data is included in the aggregation, the model may begin to abstract away modality-specific inconsistencies and instead learn robust decision boundaries that generalize across latent spaces. Nevertheless, the persistent underperformance of annotation embeddings, especially in composite settings, underscores the importance of selecting compatible embedding strategies when integrating unaligned modalities. These findings suggest that the structural properties of embedding spaces—particularly whether they support compositionality, smoothness, and alignment—play a critical role in determining model robustness and generalization in multi-modal learning frameworks.

Future work could explore alternative embedding strategies for annotations, such as fine-tuned VAEs or hybrid approaches that combine semantic and latent-space regularization, to improve consistency across modalities. Follow up research could pivot towards a heterogeneous data graph used for benchmarking. Identifying a multimodal data graph with complex relationships, structures and well defined prediction tasks will take considerable effort. This paper has demonstrated that the basic premise of multimodal embedding aggregation is viable and is able to hold information without interfering signals immediately collapsing into noise. We plan to apply this technique to encoding tumor evolution patterns, including encoded information representing tri-nucleotide patterns, structured somatic mutation timing, and subclonal mutation clustering. Other experiments could include multi-modal single cell data encoding, capturing transcriptomic and methylation data across various cell states.

## Methods

### Creation of Dataset

To obtain the necessary data, we utilized The Cancer Genome Atlas (TCGA), a comprehensive repository containing molecular and clinical data from over 10,000 cancer patients across 33 cancer types. For this study, we focused on six cancers: Breast Cancer (BRCA), Bladder Cancer (BLCA), Lung Adenocarcinoma (LUAD), Stomach Adenocarcinoma (STAD), Thyroid Cancer (THCA), and Colon Adenocarcinoma (COAD). TCGA provides extensive genomic insights, including RNA expression profiles, somatic mutations, and imaging data such as H&E-stained slides, enabling a robust analysis of cancer heterogeneity and multi modal embedding aggregations.

### Data Preparation

For each cancer type, we gathered the corresponding H&E images, patient annotations, gene mutations, and RNA readouts. Specifically, we obtained data from patients diagnosed with either Breast (BRCA), Bladder (BLCA), Lung (LUAD), Thyroid (THCA), Colon (COAD) or Stomach (STAD) cancer across these six cancer types. (Table 1, Table 2)

**Table 1:**
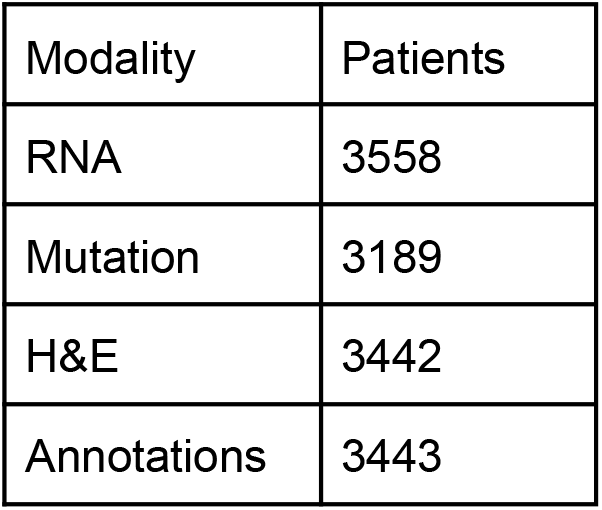
Number of patients available per data modality. RNA data includes the largest cohort with 3,558 patients, while mutation data includes the smallest cohort with 3,189 patients.

**Table 2:**
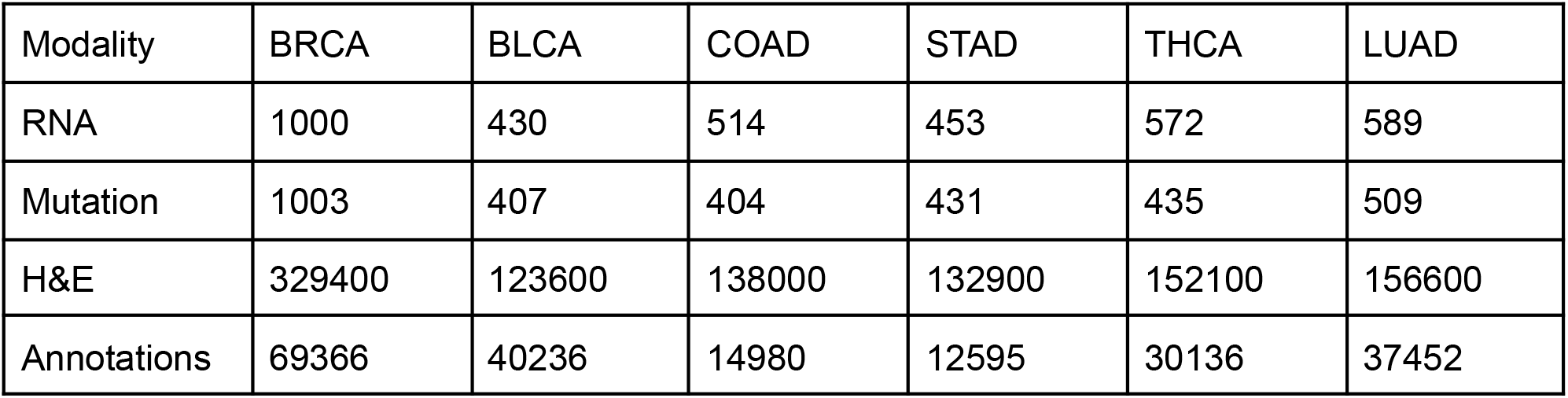
Detailed distribution of available embeddings per cancer type across data modalities. Each modality includes at least one embedding per patient. RNA and mutation modalities consistently provide one embedding per patient. In contrast, annotations and H&E image modalities often include multiple embeddings per patient, with the image modality exhibiting the widest distribution.

| Modality | BRCA | BLCA | COAD | STAD | THCA | LUAD |
| --- | --- | --- | --- | --- | --- | --- |
| RNA | 1000 | 430 | 514 | 453 | 572 | 589 |
| Mutation | 1003 | 407 | 404 | 431 | 435 | 509 |
| H&E | 329400 | 123600 | 138000 | 132900 | 152100 | 156600 |
| Annotations | 69366 | 40236 | 14980 | 12595 | 30136 | 37452 |

While image, RNA, and mutation data were readily available in formats like CSV and TIFF files, patient annotations were only provided in PDF format, making them less accessible for analysis. To facilitate processing, we converted the PDF files into text. For cases where the PDFs contained handwritten notes, we used Optical Character Recognition (OCR) to extract the textual information. The extracted text was then segmented by sentences or periods, producing multiple distinct text segments for each patient. Each segment was then transformed into separate embeddings, allowing us to create a comprehensive set of embeddings for every individual patient. This multi-layered embedding strategy ensured that all relevant data modalities were captured and integrated accurately for further analysis.

### Recognizer Network Architecture

We developed a deep learning-based recognizer network to identify the constituent modality embeddings within an aggregated embedding vector. The objective of this model was to determine whether it is possible to recover the composition of an aggregated embedding—specifically, which modality types (RNA, H&E, mutation, or annotation) were included—based solely on the vector representation. This task reflects a compositional recognition problem, where the challenge lies in disentangling heterogeneous embeddings that originate from unaligned latent spaces.

The recognizer network architecture consists of a series of shared layers followed by modality-specific output branches. The shared layers, common to all input embeddings, are designed to extract generalizable features from the aggregated vectors. These layers include standard components such as fully connected layers with activation functions, batch normalization, and dropout for regularization. After this shared feature extraction stage, the network branches into multiple modality-specific layers, each tailored to predict the presence or absence of a particular embedding type within the input vector. This modular structure allows the network to maintain generalization while preserving sensitivity to distinct modality-specific features.

To train the recognizer network, we used aggregated embeddings constructed from real patient data across four modalities: RNA-seq, H&E image embeddings, somatic mutations, and clinical annotations. The training data was generated using the modality-level sampling strategy (first strategy), in which aggregated embeddings were composed by randomly sampling modality embeddings without regard to patient identity. This setup allowed us to assess the network’s ability to learn modality-specific signatures in a mixed latent space.

We evaluated the recognizer network under two complementary experimental settings. In the simple aggregation approach, models were trained and tested on embeddings generated using fixed sample counts (isolation models), while in the composite setting, the recognizer was trained on a mixture of sample counts to assess robustness to input heterogeneity. This design enabled us to quantify the recognizer’s capacity to infer embedding composition under both controlled and variable aggregation conditions.

### Embedding Generation

To capture the unique characteristics of each modality—RNA expression, somatic mutations, histopathology (H&E) images, and patient-level annotations—we generated modality-specific embeddings designed to reside in a shared representational space. Given the inherent heterogeneity in feature types and dimensionalities across these data sources, our objective was to project each modality into a compatible latent space that retained biological relevance while enabling downstream integration.

For RNA, mutation, and image data, we employed Variational Autoencoders (VAEs) to learn compact, biologically meaningful latent representations. For patient-level annotations, we leveraged a pretrained Sentence-BERT (sBERT)model, which produces 768-dimensional embeddings optimized for semantic similarity in textual data (https://sbert.net/). To align all modality representations, each VAE was configured to output embeddings of 768 dimensions, matching the fixed dimensionality of sBERT. This uniform representation allowed for straightforward aggregation and cross-modality comparison.

Each VAE was trained separately on data from all six cancer types to ensure generalizability and to preserve the biological variability inherent across different tumor types. After training, embeddings were extracted using the encoder component of each VAE, enabling compression of modality-specific input into a biologically informative latent space.

For somatic mutations, the mutation VAE (NETVAE) was trained on a one-hot encoded mutation matrix. Training was performed with a batch size of 256 and incorporated an early stopping criterion with a patience of 10 epochs to prevent overfitting. A maximum of 50 training epochs was allowed. The resulting encoder produced a 768-dimensional latent vector that effectively captured mutational patterns across patients.

For RNA expression data, a dedicated VAE was trained using RNA-seq data from The Cancer Genome Atlas (TCGA). Training was performed in two stages: an initial pretraining phase followed by fine-tuning. A warm-up strategy was applied to the Kullback–Leibler (KL) divergence term in the VAE loss function to improve training stability. Specifically, KL loss was initialized at zero and gradually increased according to an annealing schedule governed by a κ (kappa) parameter. In our implementation, κ was set to 1, and the KL loss weight (β) was initialized at 0, resulting in a one-epoch warm-up phase. This annealing technique is known to promote more stable convergence by allowing the model to learn meaningful reconstructions before imposing latent space regularization.

For H&E histopathology images, whole-slide images were partitioned into non-overlapping 256×256 pixel tiles. Tiles predominantly containing background were removed using a simple RGB pixel thresholding heuristic. Remaining tiles were processed with Prov-Gigapath, a whole-slide foundation model pre-trained on large-scale pathology datasets (https://github.com/prov-gigapath/prov-gigapath). Tile-level inference produced 1536-dimensional embeddings, which were then truncated to the final 768 dimensions. This truncation strategy was informed by internal benchmarking, which indicated that the latter half of the embedding vector retained sufficient discriminative power for downstream tasks. The resulting tile-level embeddings were used in all subsequent analyses.

### Embedding Aggregation

To construct a comprehensive dataset for training and evaluating recognition models, we generated aggregated embeddings composed of three to ten constituent embeddings per vector. This range ensured exposure to varying levels of aggregation complexity, enabling the model to generalize across different embedding combinations. The embedding construction process followed two distinct sampling strategies: the simple aggregation approach and the specific-cancer approach, corresponding to our previously described first and second strategies, respectively.

For the simple aggregation approach, all available embeddings across modalities—RNA, H&E, mutations, and annotations—were first loaded into memory. A random sampling procedure was applied, selecting a specified number of embeddings (e.g., sample count of 3 to 10) regardless of their patient or cancer type of origin. The selected embeddings were summed to produce a new 768-dimensional aggregated embedding, and the modalities contributing to the aggregation were tracked and stored as ground truth labels for supervised training.

In the specific-cancer approach, the sampling procedure was constrained to enforce cancer-type specificity. After loading all available embeddings, random sampling was performed such that all selected embeddings originated from the same cancer type (e.g., only embeddings derived from BRCA patients). As in the simple approach, the selected embeddings were summed to form a 768-dimensional vector. Both the modalities involved in the aggregation and the corresponding cancer type were recorded as ground truth labels. This design enabled us to evaluate the recognizer model’s performance under both general and cancer-specific conditions, while preserving control over the composition and structure of the aggregated embeddings.

### Training the Recognizer Network

To evaluate whether the composition of aggregated embeddings could be recovered, we trained a deep learning-based recognizer network on vectors composed of three to ten constituent embeddings. These embeddings were generated through the simple aggregation approach, with additional models trained under cancer-type-specific sampling to assess performance under more biologically constrained conditions.

Aggregated embeddings were constructed by randomly selecting and summing modality-specific embeddings drawn from RNA expression, H&E images, somatic mutations, and patient-level annotations. This procedure produced 768-dimensional vectors, each labeled with the set of contributing modalities and, when applicable, the associated cancer type. The resulting dataset formed the input for training isolation recognizer models, each trained on a specific sample count, as well as a composite recognizer model, trained on the full range of sample counts (3–10) within a unified dataset.

### Training Procedure

The recognizer network was trained using supervised learning. The input to the model was the aggregated embedding vector, and the output was a binary vector indicating the presence or absence of each modality in the aggregation. The model was trained using the Adam optimizer with an initial learning rate of 0.001, and the mean squared error (MSE) loss function was used to penalize incorrect predictions of modality presence.

The network architecture consisted of an initial set of shared layers designed for general feature extraction, followed by modality-specific output branches that performed binary classification for each of the four modalities. To improve generalization and stability, the architecture incorporated fully connected layers, dropout, and batch normalization.

To prevent modality imbalance during training, we explicitly controlled the probability of selecting each modality during the random sampling process. For example, when using four modalities, each was assigned a uniform selection probability of 25%, ensuring equal representation across training batches and avoiding bias toward more frequently represented modalities.

### Validation and Testing

The dataset was split into training, validation, and test sets using an 80/20 split, followed by an 80/20 subdivision of the training set into training and validation subsets. The validation set was used during training to monitor model generalization and tune hyperparameters, while the test set was held out entirely for final performance evaluation.

### Composite Recognizer Model

In addition to training individual isolation models for each sample count, we trained a composite recognizer modelusing a merged dataset containing all aggregated embeddings with sample counts from 3 to 10. This model was designed to assess whether a single network could generalize across varying levels of aggregation complexity and modality combinations. The same training architecture and optimization parameters were applied to the composite model, enabling direct comparison with the isolation-based models.

### Metric calculation for recognizer network evaluation

The evaluation of the recognizer network required special consideration due to the sparse structure of the training data, which was generated using the first sampling strategy—modality-level sampling. In this approach, embeddings were randomly selected across modalities without regard to patient identity, and aggregated to create synthetic vectors composed of three to ten constituent embeddings. Because any given modality may or may not have been included in a given aggregated vector, the resulting label matrix contained a high proportion of zeros. Additionally, constraints in the data generation process (e.g., limiting the number of embeddings or excluding certain modalities) led to systematically enforced zero entries in specific columns.

This sparsity posed challenges for standard classification metrics such as accuracy, F1 score, and Matthews correlation coefficient (MCC). A trivial model that always predicts absence (i.e., zeros for all modality labels) could achieve high accuracy, especially in test sets with many modality-absent samples. To ensure correct and informative evaluation, we stratified metric computation for each modality (RNA, H&E, somatic mutations, and clinical annotations) by separating test samples into two groups:

1. Zero-labeled samples, in which a modality was absent from the aggregated vector
2. Non-zero-labeled samples, in which a modality was present and thus had to be correctly recognized

Performance metrics—MCC, accuracy, precision, recall, and F1 score—were computed independently within each group. This allowed us to assess the recognizer model’s capacity to both correctly detect present modalities and avoid false positives for absent ones, preventing inflated performance due to label imbalance.

For models trained under noise-perturbed conditions, we applied the same stratification logic. These datasets were derived from clean aggregated vectors (generated via modality-level sampling) with Gaussian noise introduced to replace a subset of embeddings. While the total number of embeddings per vector remained unchanged, the substitution of valid modality embeddings with noise created further sparsity in the ground truth labels. In this setting, models predicting only absent modalities could again appear to perform well based on raw accuracy, despite lacking real compositional understanding. As with the standard setting, excluding zero-only rows and stratifying metrics enabled a more meaningful evaluation of the model’s robustness.

By applying these adjustments, we ensured that the reported metrics provided a reliable assessment of the recognizer network’s performance in identifying the constituent modality composition of aggregated embeddings, both under standard conditions and in the presence of noise.

### Gaussian Mixture Models

To enhance the visualization of information retrieval, we employed Gaussian Mixture Models (GMMs). GMMs are probabilistic models that represent complex data distributions as a combination of multiple Gaussian components, each weighted to capture distinct patterns or clusters. This approach is widely used in unsupervised learning for clustering and anomaly detection. In our analysis, we constructed a GMM with six components to evaluate its ability to distinguish between the six cancer types in our dataset. The primary goal was to compare a 2D principal component analysis (PCA) visualization with GMM clustering to assess their alignment. While biological non-linearity prevents a clear-cut separation between cancer types, distinct cluster boundaries emerged, highlighting the potential of GMMs in capturing underlying structures in high-dimensional biological data.

### Classification Model

The classification model was trained using a supervised learning approach, with input features consisting of aggregated and concatenated patient-level embeddings constructed via the second sampling strategy (patient-level sampling). Each embedding was paired with a ground truth label corresponding to one of three classification targets: cancer type, cancer subtype, or tumor mutational burden (TMB).

Data were split using an 80/20 train-test split, with training and testing set ratios held consistent across all experimental runs. To ensure reliable and robust performance estimation, the model was trained and evaluated across at least 30 independent runs. This repeated training procedure reduced the impact of stochastic variability introduced by random initialization and data shuffling, and enabled the calculation of stable performance metrics across runs. Results were aggregated to report average performance, allowing for a more accurate and reproducible assessment of the model’s predictive capacity.

### Tumor Mutational Burden (TMB) Calculation

To calculate the tumor mutational burden (TMB) for each patient, we used the one-hot encoded somatic mutation file, which was used to generate the mutation embeddings, where each gene’s mutation status was encoded as either mutated (1) or not mutated (0). For each patient, we counted the number of mutated genes by summing all values across the remaining columns.

To standardize the calculation, we assumed an exonic coverage of 30 megabases (Mb) and computed TMB as the number of mutations per megabase using this formula: *TMB = Number of Mutations / 30. [21]*

We then classified TMB into high and low categories based on a predefined threshold. A TMB value of 0.5 mutations/Mb or higher was classified as high (1), while values below this threshold were classified as low (0).

If no mutations were detected for a patient, TMB was set to 0, and the classification was assigned a separate category (2) to account for cases where no mutations were present.

This approach ensures a consistent and interpretable quantification of tumor mutational burden across patients while enabling classification into biologically relevant groups.

### Tumor Subtypes

We utilized publicly available resources to map patients and cancer types to their corresponding cancer subtypes. Specifically, subtype annotations were derived using data and tools from the Genomic Data Commons (GDC) [22] and the subtype mapping file provided in the associated repository (https://github.com/NCICCGPO/gdan-tmp-models/blob/main/tools/cancer2name.json). Subtype classification was performed only for cancer types where a sufficient number of annotated samples (n > 100) were available to support reliable supervised learning.

## Acknowledgements

The research supporting this publication was supported by NCI GDAN 5U24CA264007, NIH U54HG012517 and NHGRI U24HG010263. The research reported in this publication used computational infrastructure supported by the Office of Research Infrastructure Programs, Office of the Director, of the National Institutes of Health under Award Number S10OD034224. The content is solely the responsibility of the authors and does not necessarily represent the official views of the National Institutes of Health.

