## Supplemental Figures for "Aggregating multimodal cancer data across unaligned embedding spaces maintains tumor of origin signal"

SC: 3 R: 3

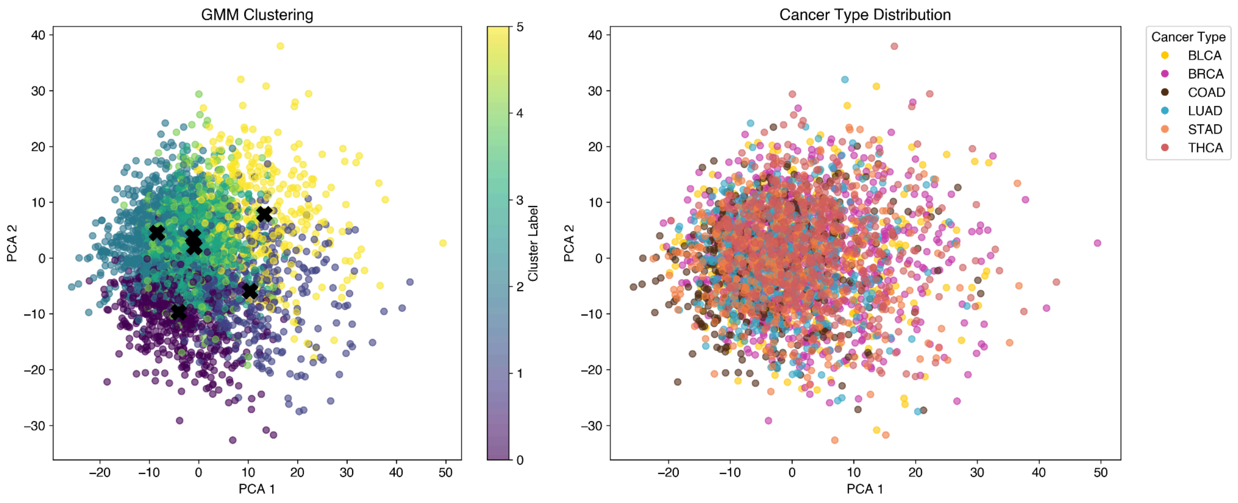


SC: 3 R: 4

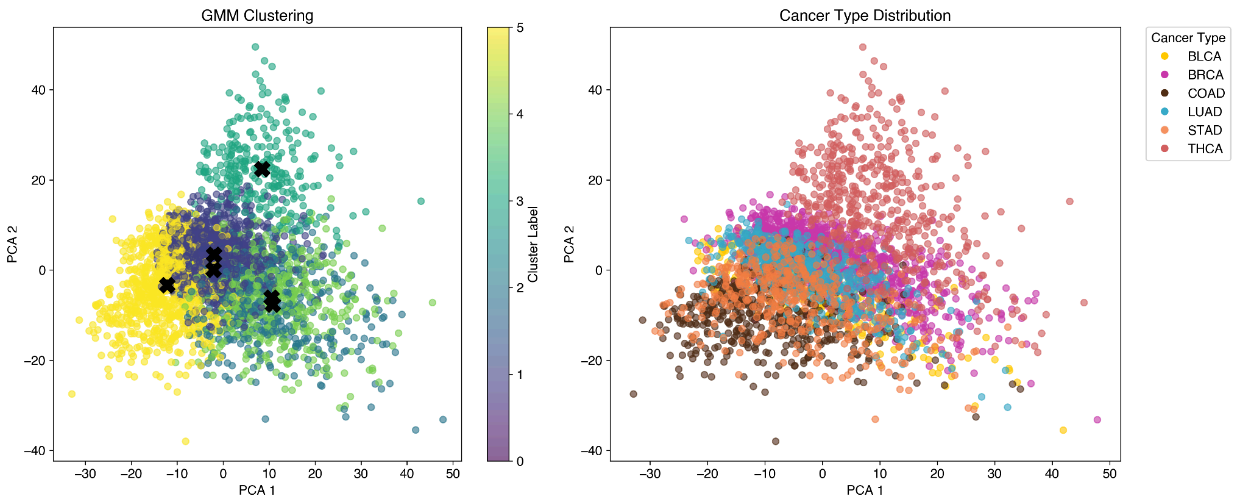


SC: 3 R: 5

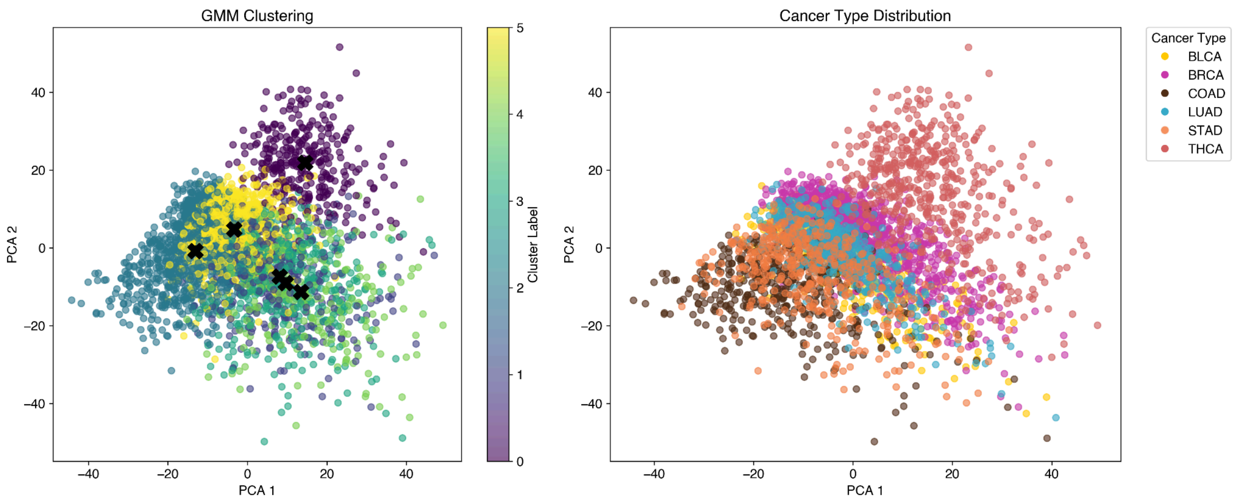

**Supplemental Figure 1**: Clustering performance with a fixed sample count of 3 and increasing sampling counts from 3 to 5 demonstrates progressive enhancement in cluster separation. With 3 repetitions, clustering lacks clear separation, as reflected by the Gaussian Mixture Model (GMM) clustering. Increasing the repetition count improves both cancer type and GMM clustering, resulting in greater separation and more distinct cluster formation.

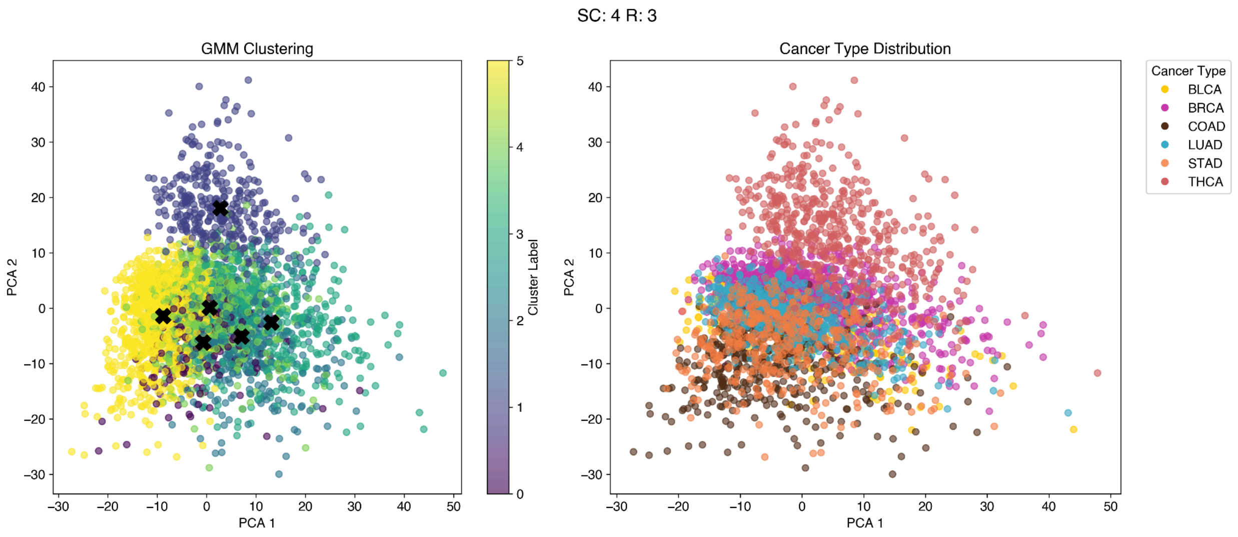
SC: 4 R: 3
GMM Clustering
PCA 1
PCA 2
Cluster Label
Cancer Type Distribution
Cancer Type
BLCA
BRCA
COAD
LUAD
STAD
THCA

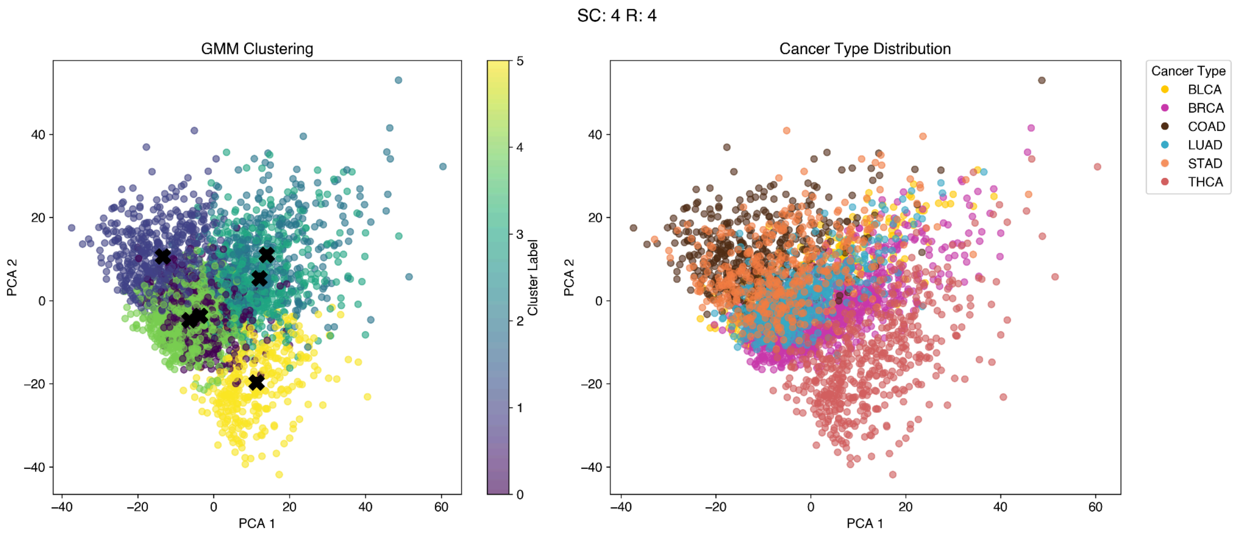
SC: 4 R: 4
GMM Clustering
PCA 1
PCA 2
Cluster Label
Cancer Type Distribution
Cancer Type
BLCA
BRCA
COAD
LUAD
STAD
THCA

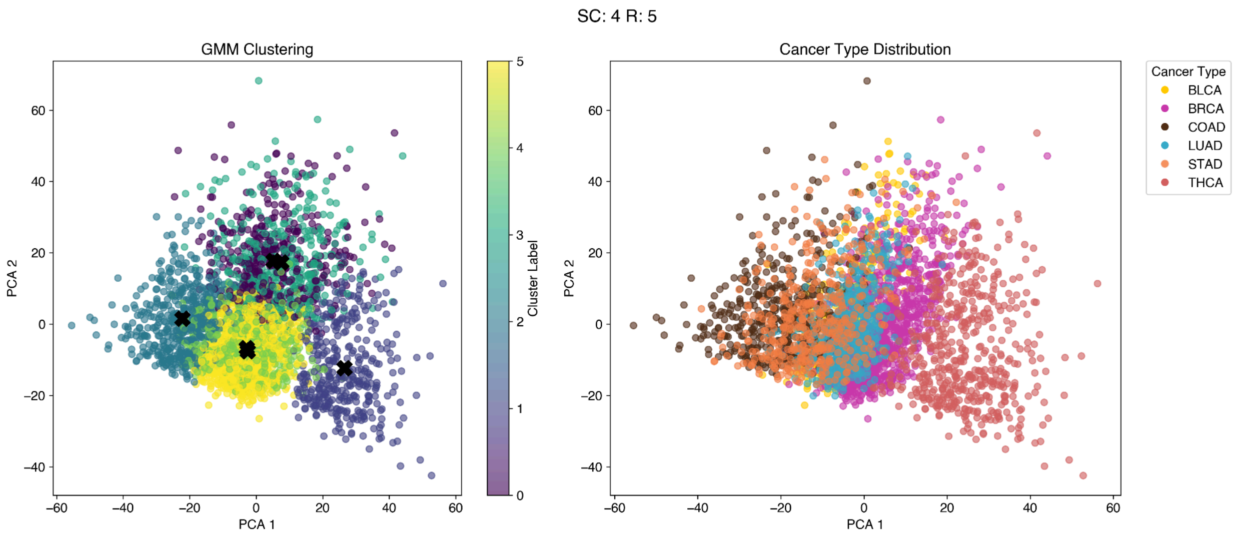
SC: 4 R: 5
GMM Clustering
PCA 1
PCA 2
Cluster Label
Cancer Type Distribution
Cancer Type
BLCA
BRCA
COAD
LUAD
STAD
THCA

**Supplemental Figure 2**: Clustering performance with a fixed sample count of 4 and varying repetition counts between 3 and 5 shows enhanced clustering quality with increasing repetition. As repetition count increases, both Gaussian Mixture Model (GMM) clustering and cancer type clustering exhibit improved alignment and separation, indicating more robust cluster formation.

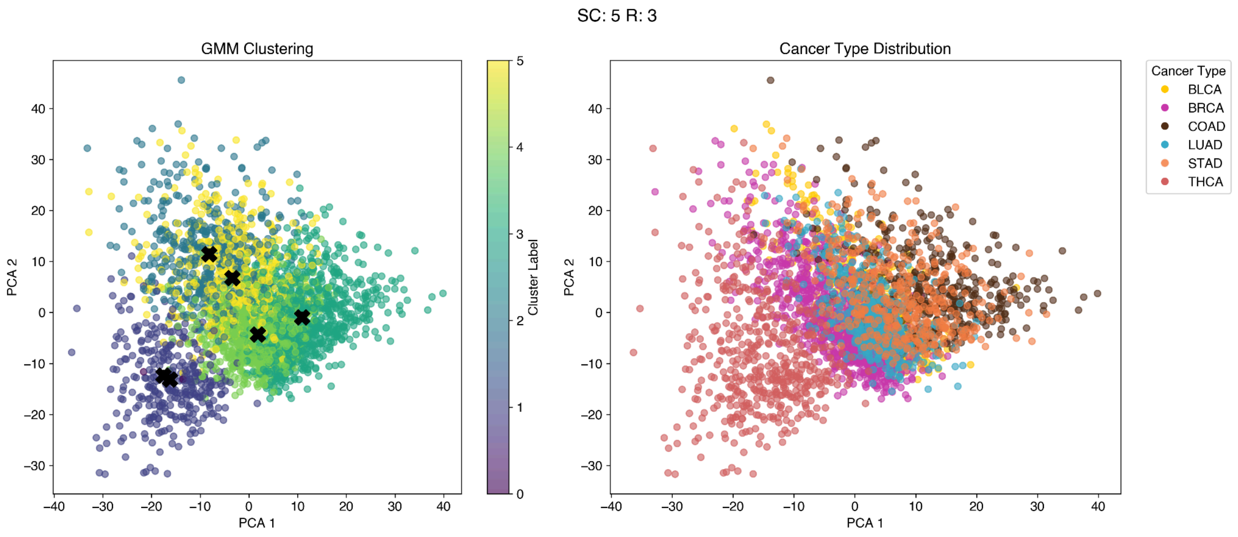
SC: 5 R: 3
GMM Clustering
PCA 1
PCA 2
Cluster Label
Cancer Type Distribution
Cancer Type
BLCA
BRCA
COAD
LUAD
STAD
THCA

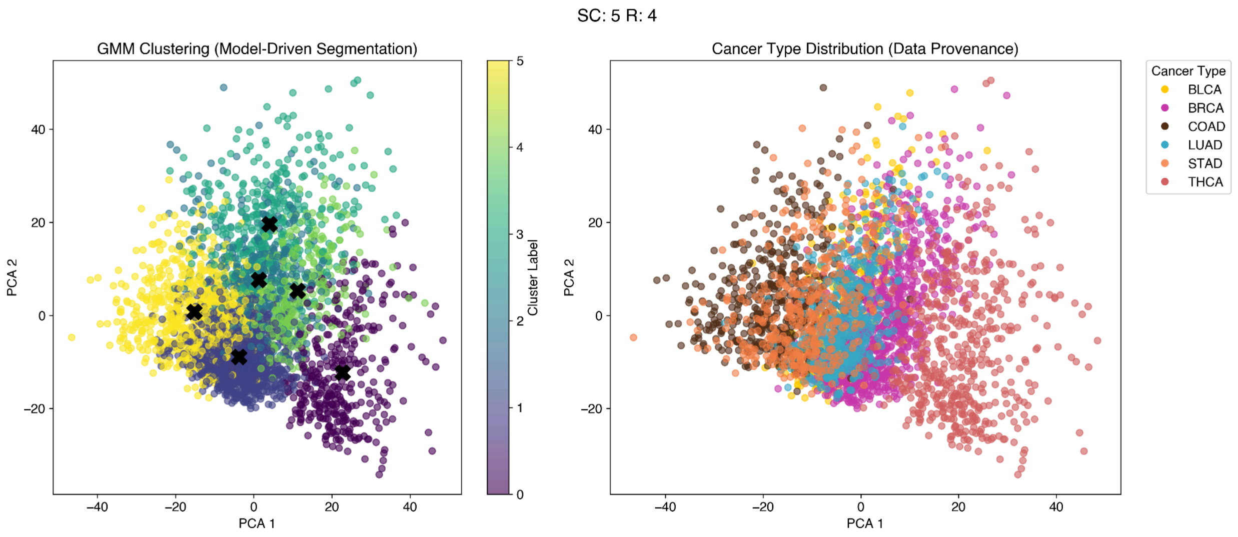
SC: 5 R: 4
GMM Clustering (Model-Driven Segmentation)
PCA 1
PCA 2
Cluster Label
Cancer Type Distribution (Data Provenance)
Cancer Type
BLCA
BRCA
COAD
LUAD
STAD
THCA

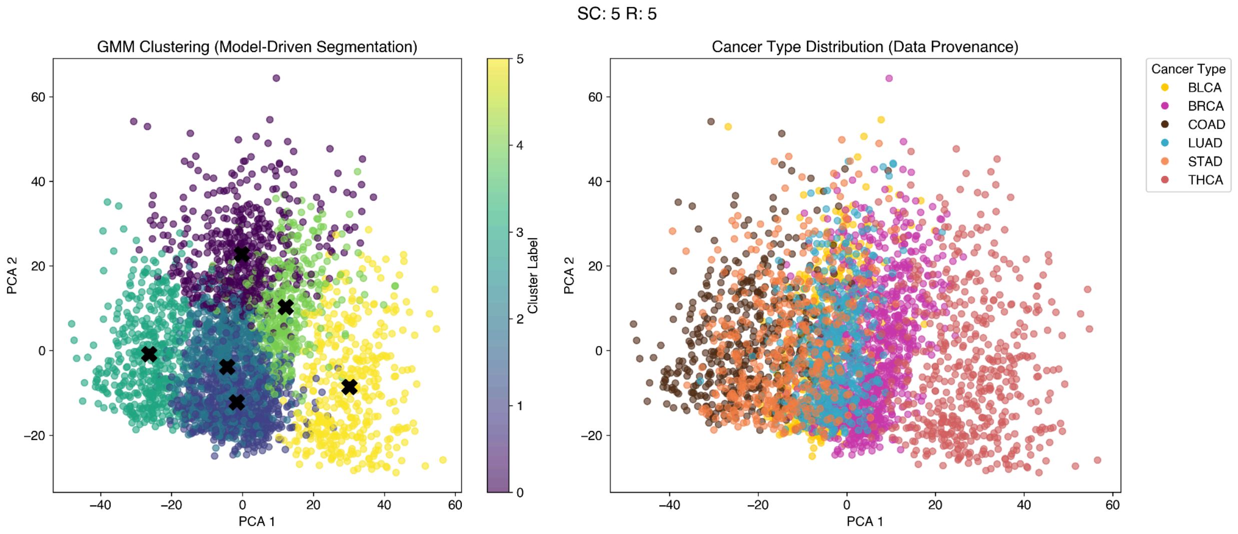
SC: 5 R: 5
GMM Clustering (Model-Driven Segmentation)
PCA 1
PCA 2
Cluster Label
Cancer Type Distribution (Data Provenance)
Cancer Type
BLCA
BRCA
COAD
LUAD
STAD
THCA

**Supplemental Figure 3**: Clustering performance with a fixed sample count of 5 and varying repetition counts between 3 and 5 demonstrates improved clustering with increasing repetition. Additionally, the Gaussian Mixture Model (GMM) unsupervised clustering aligns well with the cancer subtype clusters, indicating strong agreement between predicted and true cluster structures.

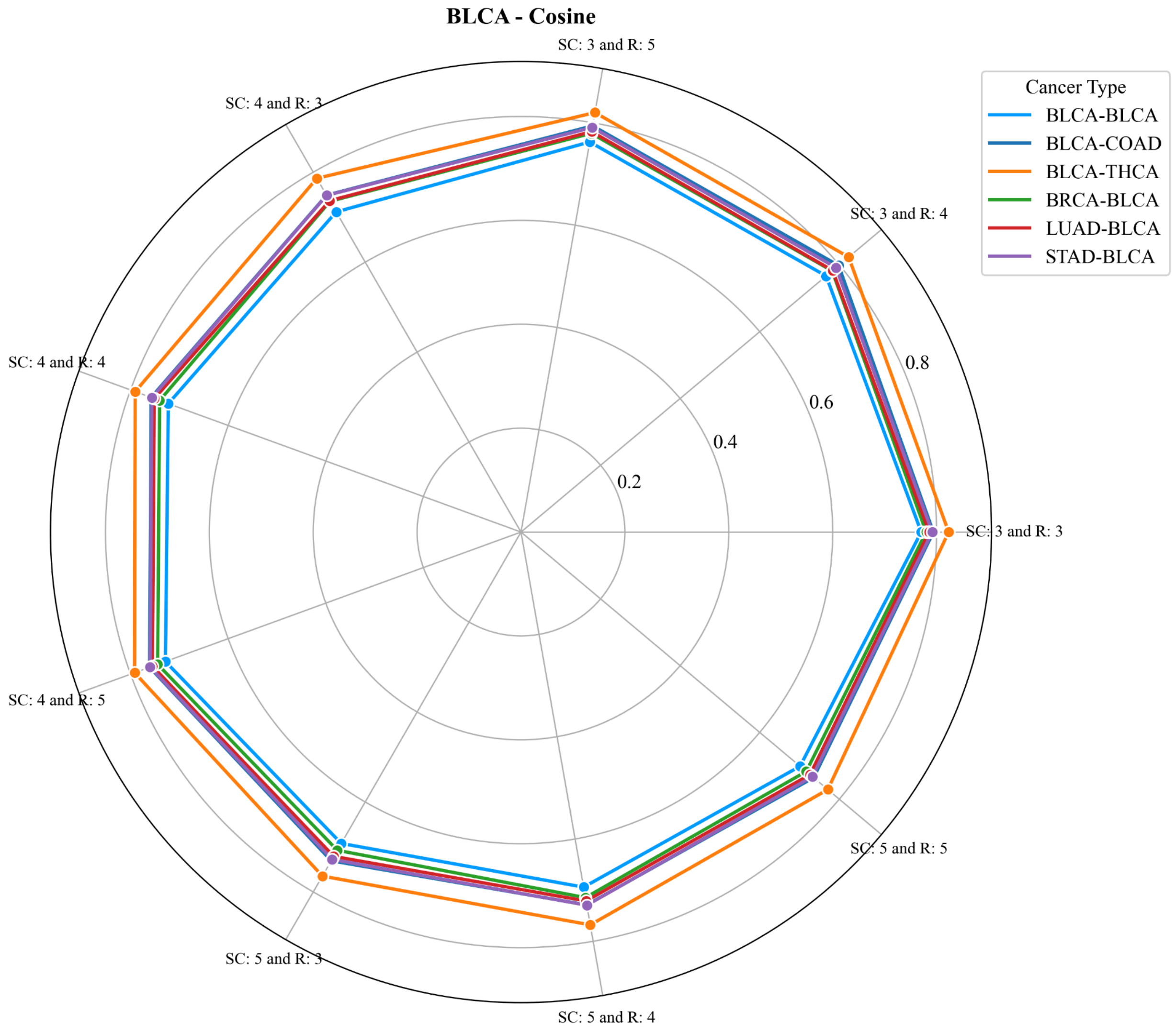
BLCA - Cosine
SC: 3 and R: 5
SC: 3 and R: 4
SC: 3 and R: 3
SC: 5 and R: 5
SC: 5 and R: 4
SC: 5 and R: 3
SC: 4 and R: 5
SC: 4 and R: 4
SC: 4 and R: 3
0.2
0.4
0.6
0.8
Cancer Type
BLCA-BLCA
BLCA-COAD
BLCA-THCA
BRCA-BLCA
LUAD-BLCA
STAD-BLCA

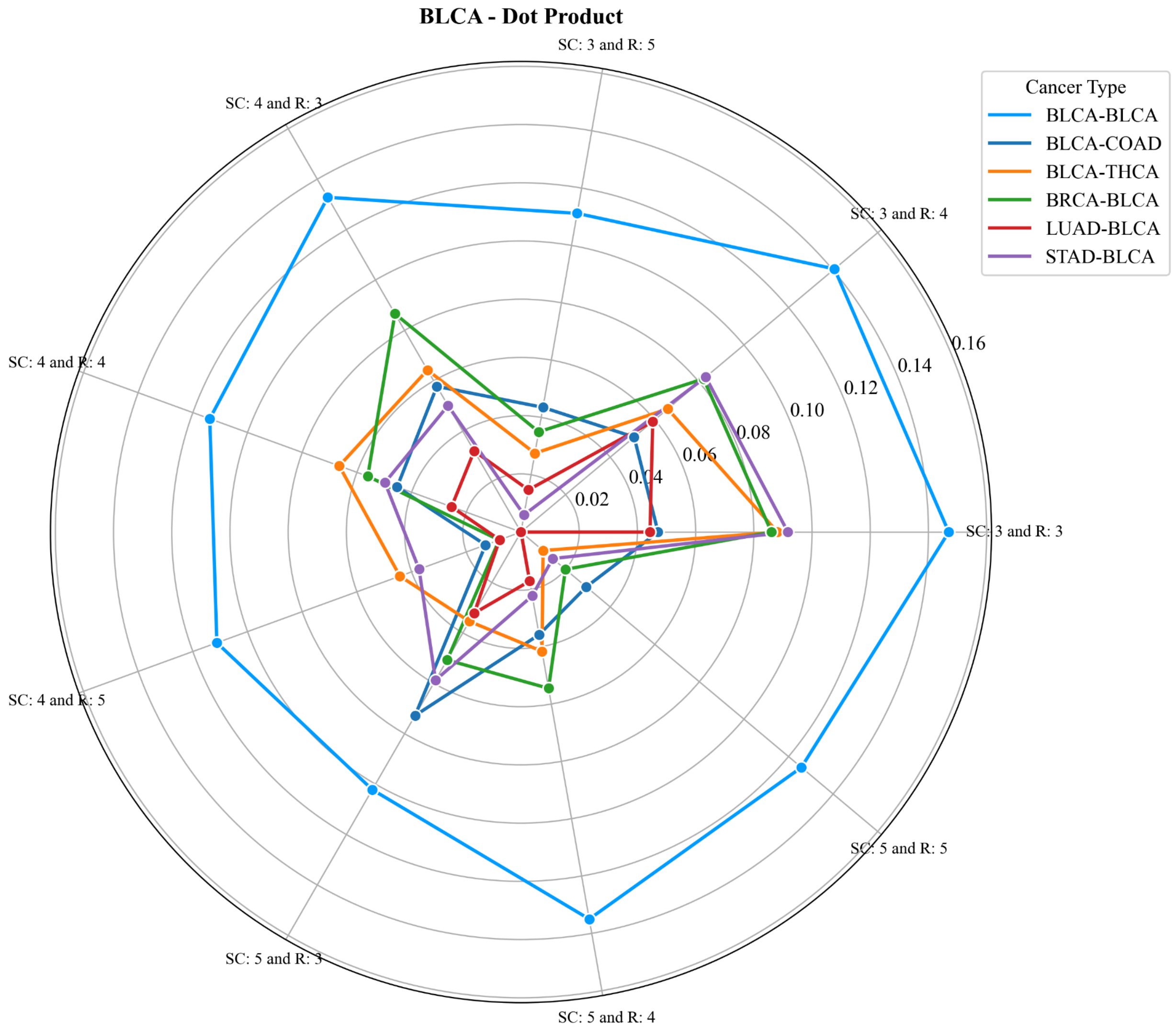

BLCA - Dot Product
SC: 3 and R: 3
SC: 3 and R: 4
SC: 3 and R: 5
SC: 4 and R: 3
SC: 4 and R: 4
SC: 4 and R: 5
SC: 5 and R: 3
SC: 5 and R: 4
SC: 5 and R: 5
0.02
0.04
0.06
0.08
0.10
0.12
0.14
0.16
Cancer Type
BLCA-BLCA
BLCA-COAD
BLCA-THCA
BRCA-BLCA
LUAD-BLCA
STAD-BLCA

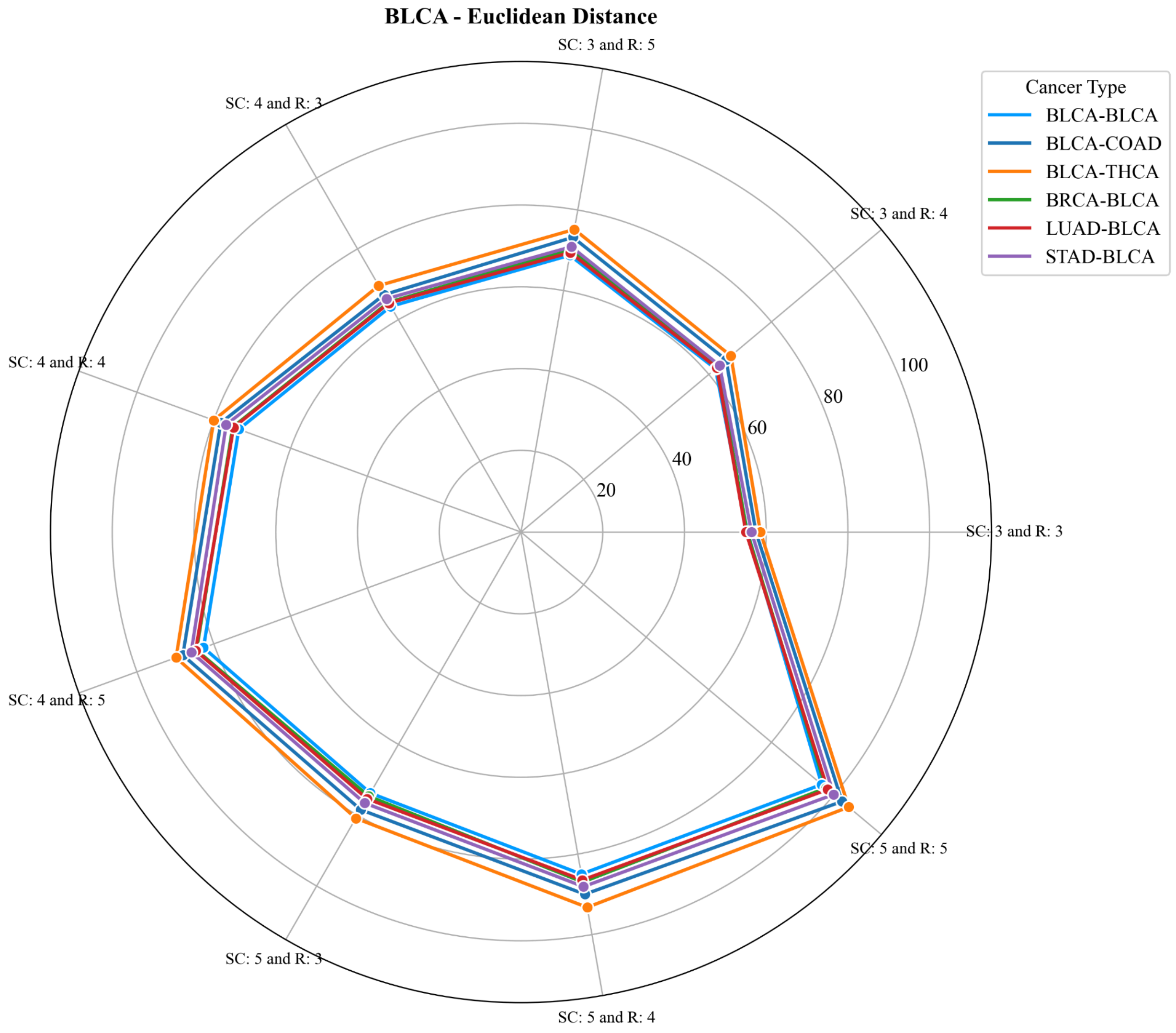
BLCA - Euclidean Distance
SC: 3 and R: 5
SC: 4 and R: 3
SC: 3 and R: 4
SC: 4 and R: 4
SC: 3 and R: 3
SC: 4 and R: 5
SC: 5 and R: 5
SC: 5 and R: 3
SC: 5 and R: 4
20
40
60
80
100
Cancer Type
BLCA-BLCA
BLCA-COAD
BLCA-THCA
BRCA-BLCA
LUAD-BLCA
STAD-BLCA

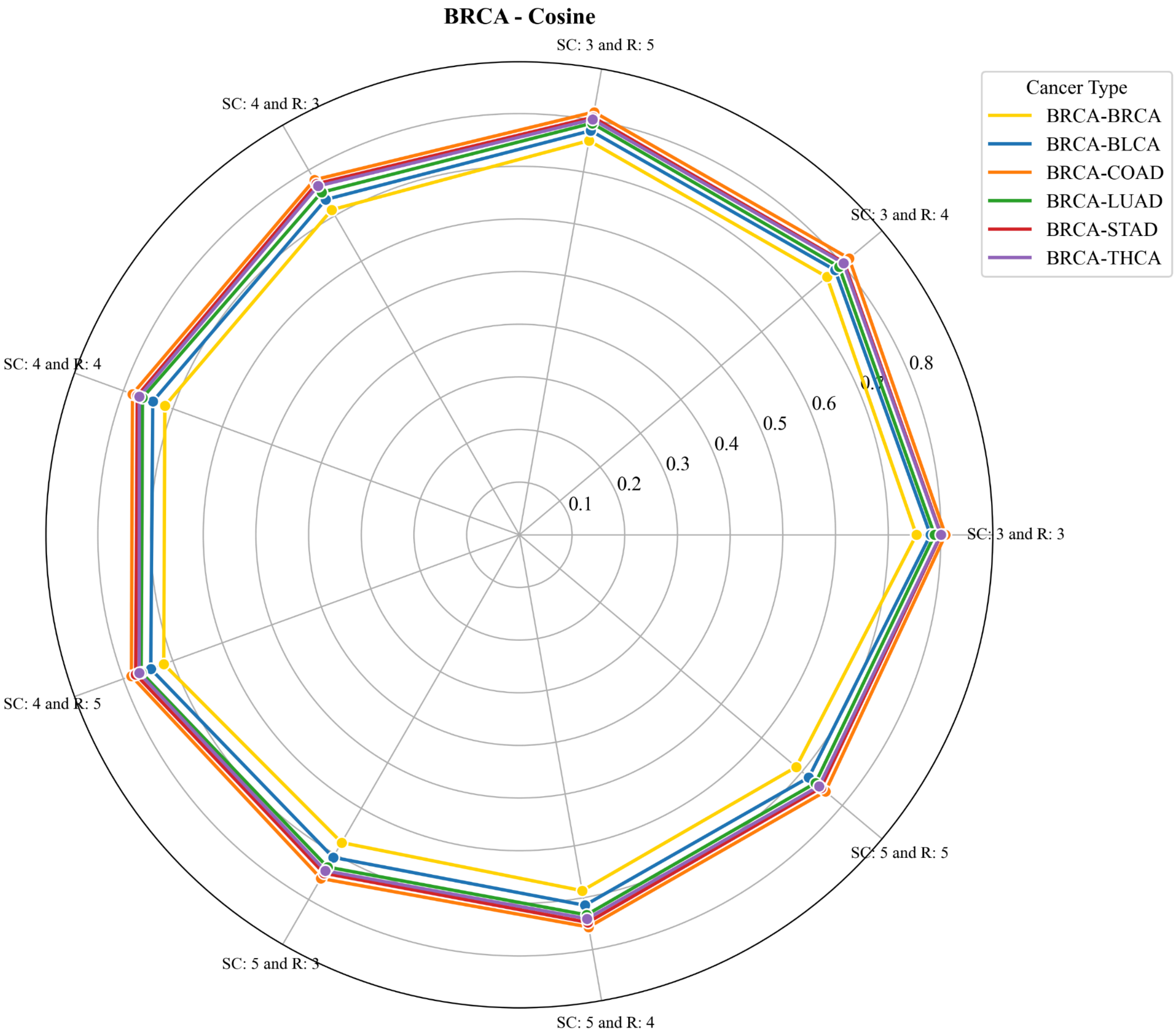

BRCA - Cosine
SC: 3 and R: 5
SC: 3 and R: 4
SC: 3 and R: 3
SC: 5 and R: 5
SC: 5 and R: 4
SC: 5 and R: 3
SC: 4 and R: 5
SC: 4 and R: 4
SC: 4 and R: 3
0.1
0.2
0.3
0.4
0.5
0.6
0.7
0.8
Cancer Type
BRCA-BRCA
BRCA-BLCA
BRCA-COAD
BRCA-LUAD
BRCA-STAD
BRCA-THCA

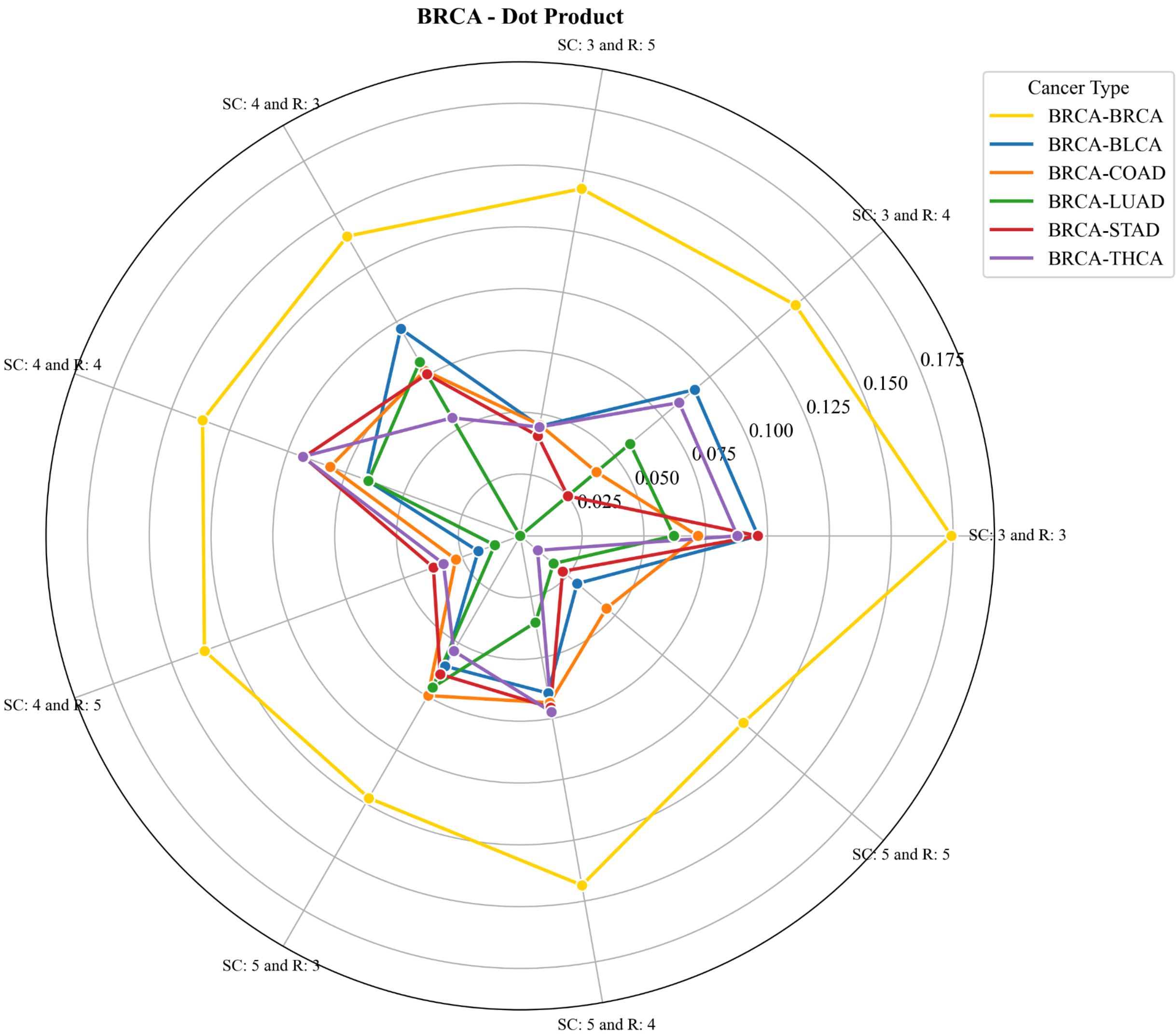
BRCA - Dot Product
SC: 3 and R: 5
SC: 3 and R: 4
SC: 3 and R: 3
SC: 5 and R: 5
SC: 5 and R: 4
SC: 5 and R: 3
SC: 4 and R: 5
SC: 4 and R: 4
SC: 4 and R: 3
0.025
0.050
0.075
0.100
0.125
0.150
0.175
Cancer Type
BRCA-BRCA
BRCA-BLCA
BRCA-COAD
BRCA-LUAD
BRCA-STAD
BRCA-THCA

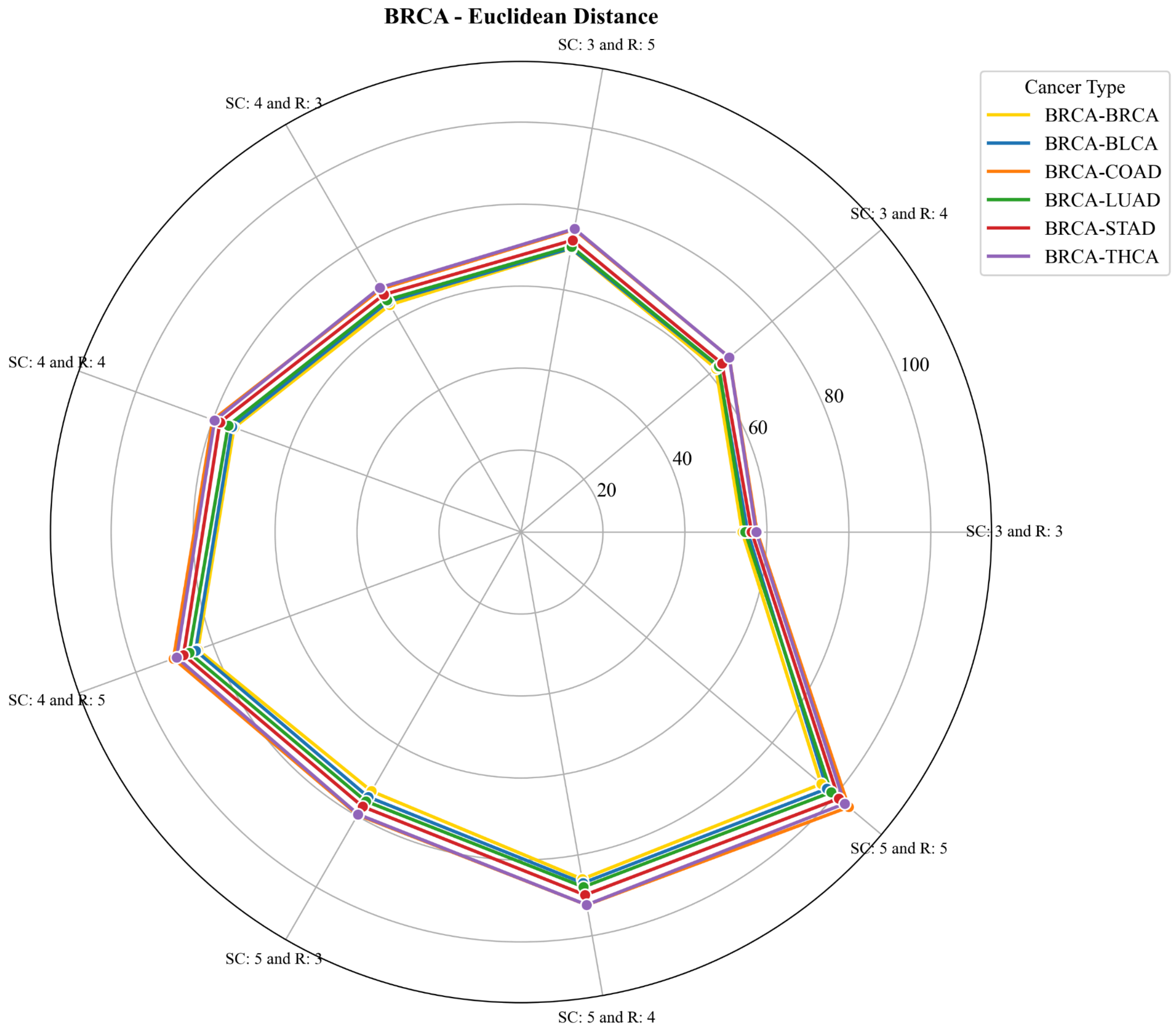

BRCA - Euclidean Distance
SC: 3 and R: 5
SC: 3 and R: 4
SC: 3 and R: 3
SC: 5 and R: 5
SC: 5 and R: 4
SC: 5 and R: 3
SC: 4 and R: 5
SC: 4 and R: 4
SC: 4 and R: 3
20
40
60
80
100
Cancer Type
BRCA-BRCA
BRCA-BLCA
BRCA-COAD
BRCA-LUAD
BRCA-STAD
BRCA-THCA

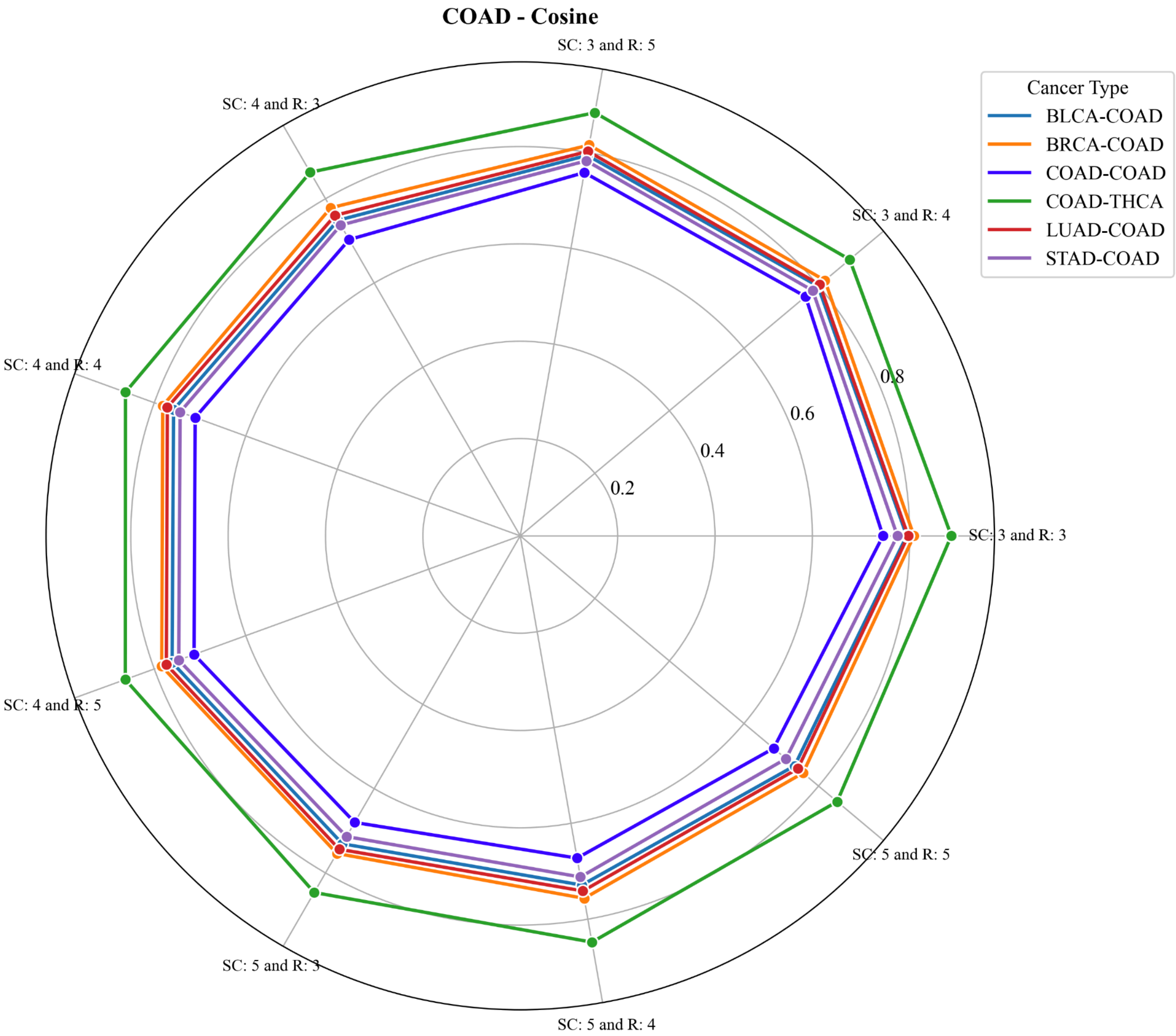
COAD - Cosine
SC: 3 and R: 5
SC: 3 and R: 4
SC: 3 and R: 3
SC: 5 and R: 5
SC: 5 and R: 4
SC: 5 and R: 3
SC: 4 and R: 5
SC: 4 and R: 4
SC: 4 and R: 3
0.2
0.4
0.6
0.8
Cancer Type
BLCA-COAD
BRCA-COAD
COAD-COAD
COAD-THCA
LUAD-COAD
STAD-COAD

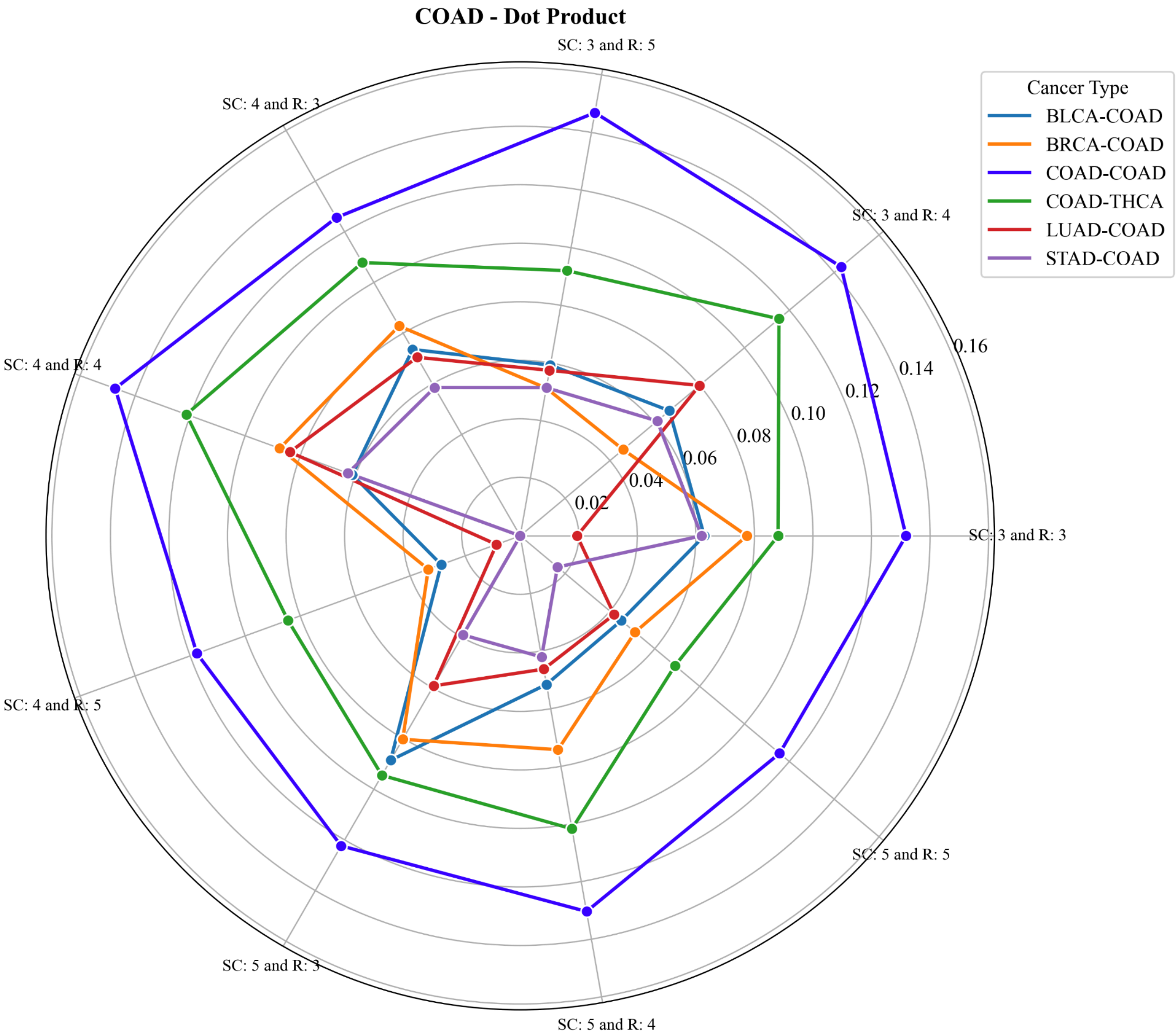

COAD - Dot Product
SC: 3 and R: 5
SC: 3 and R: 4
SC: 3 and R: 3
SC: 5 and R: 5
SC: 5 and R: 4
SC: 5 and R: 3
SC: 4 and R: 5
SC: 4 and R: 4
SC: 4 and R: 3
0.02
0.04
0.06
0.08
0.10
0.12
0.14
0.16
Cancer Type
BLCA-COAD
BRCA-COAD
COAD-COAD
COAD-THCA
LUAD-COAD
STAD-COAD

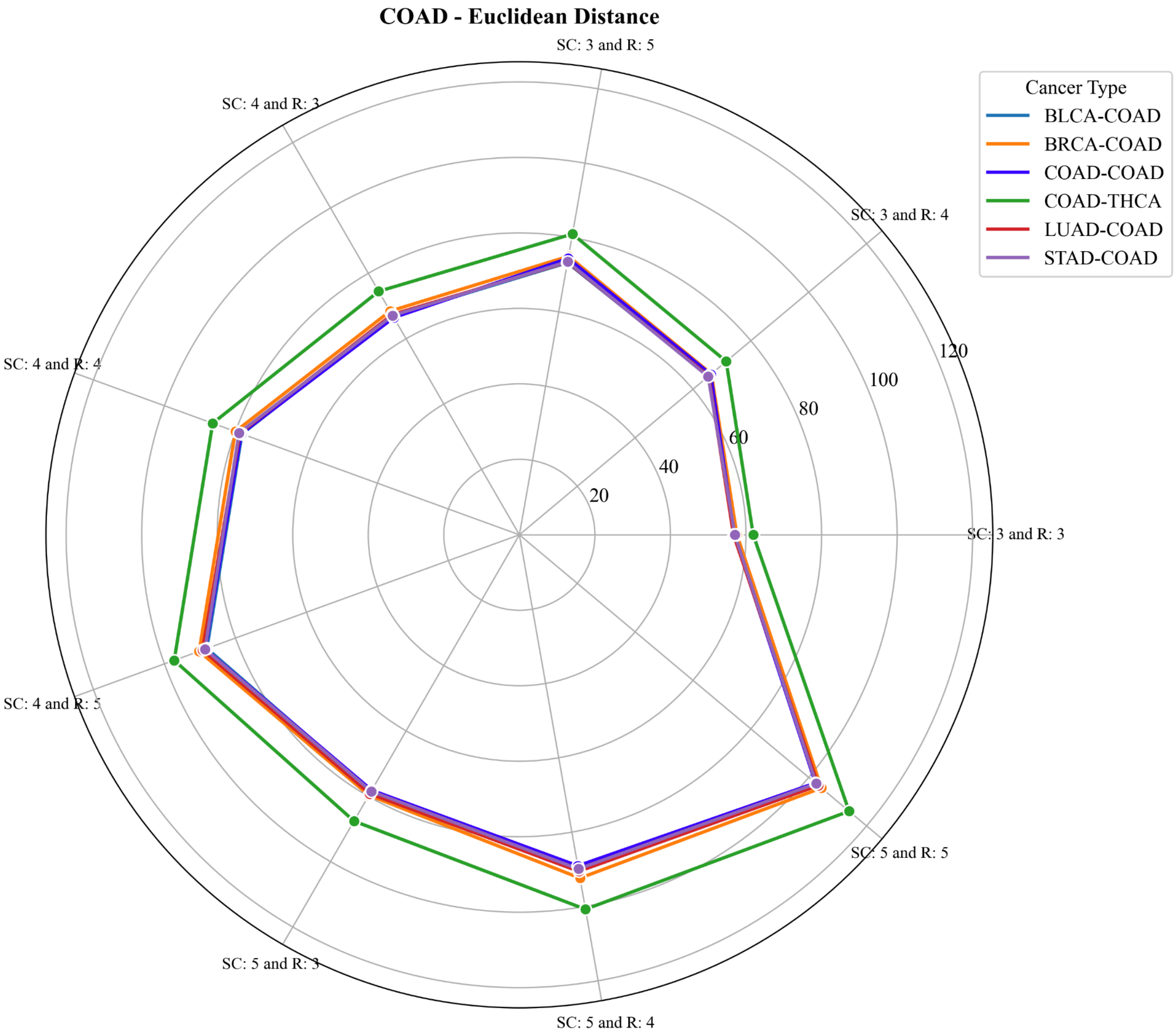
COAD - Euclidean Distance
SC: 3 and R: 5
SC: 3 and R: 4
SC: 3 and R: 3
SC: 5 and R: 5
SC: 5 and R: 4
SC: 5 and R: 3
SC: 4 and R: 5
SC: 4 and R: 4
SC: 4 and R: 3
20
40
60
80
100
120
Cancer Type
BLCA-COAD
BRCA-COAD
COAD-COAD
COAD-THCA
LUAD-COAD
STAD-COAD

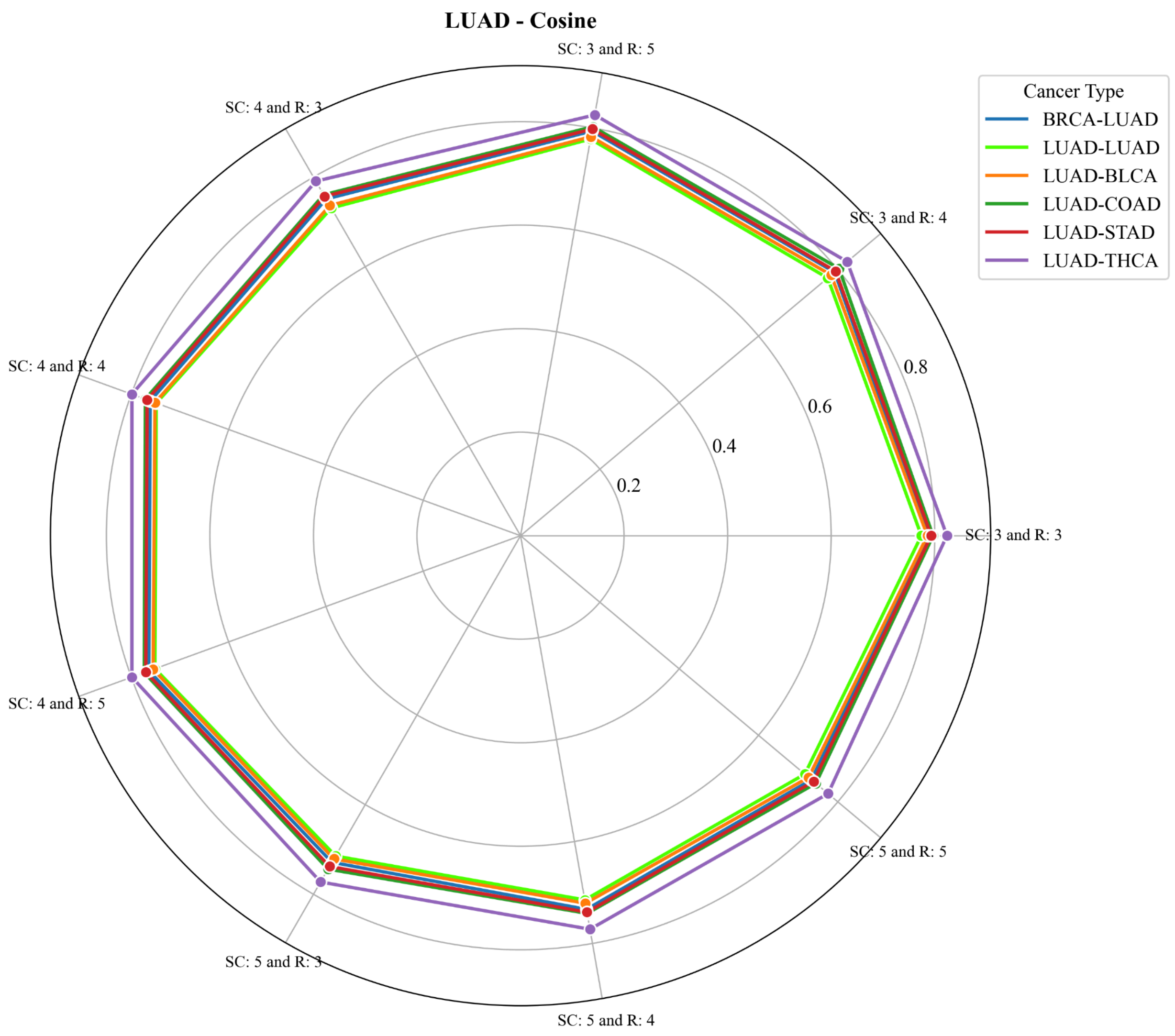

LUAD - Cosine
SC: 3 and R: 5
SC: 3 and R: 4
SC: 3 and R: 3
SC: 5 and R: 5
SC: 5 and R: 4
SC: 5 and R: 3
SC: 4 and R: 5
SC: 4 and R: 4
SC: 4 and R: 3
0.2
0.4
0.6
0.8
Cancer Type
BRCA-LUAD
LUAD-LUAD
LUAD-BLCA
LUAD-COAD
LUAD-STAD
LUAD-THCA

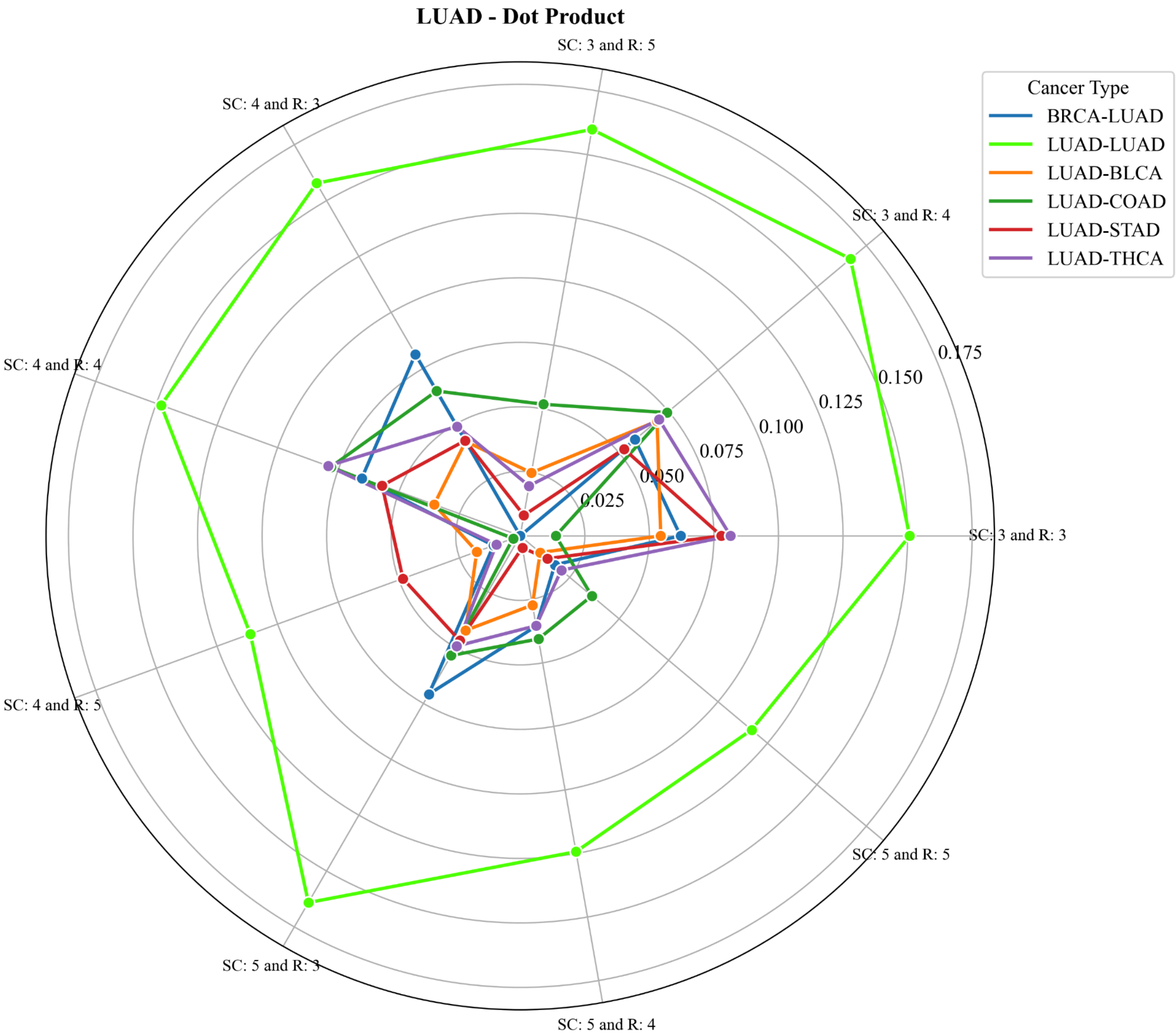
LUAD - Dot Product
SC: 3 and R: 5
SC: 3 and R: 4
SC: 3 and R: 3
SC: 5 and R: 5
SC: 5 and R: 4
SC: 5 and R: 3
SC: 4 and R: 5
SC: 4 and R: 4
SC: 4 and R: 3
0.025
0.050
0.075
0.100
0.125
0.150
0.175
Cancer Type
BRCA-LUAD
LUAD-LUAD
LUAD-BLCA
LUAD-COAD
LUAD-STAD
LUAD-THCA

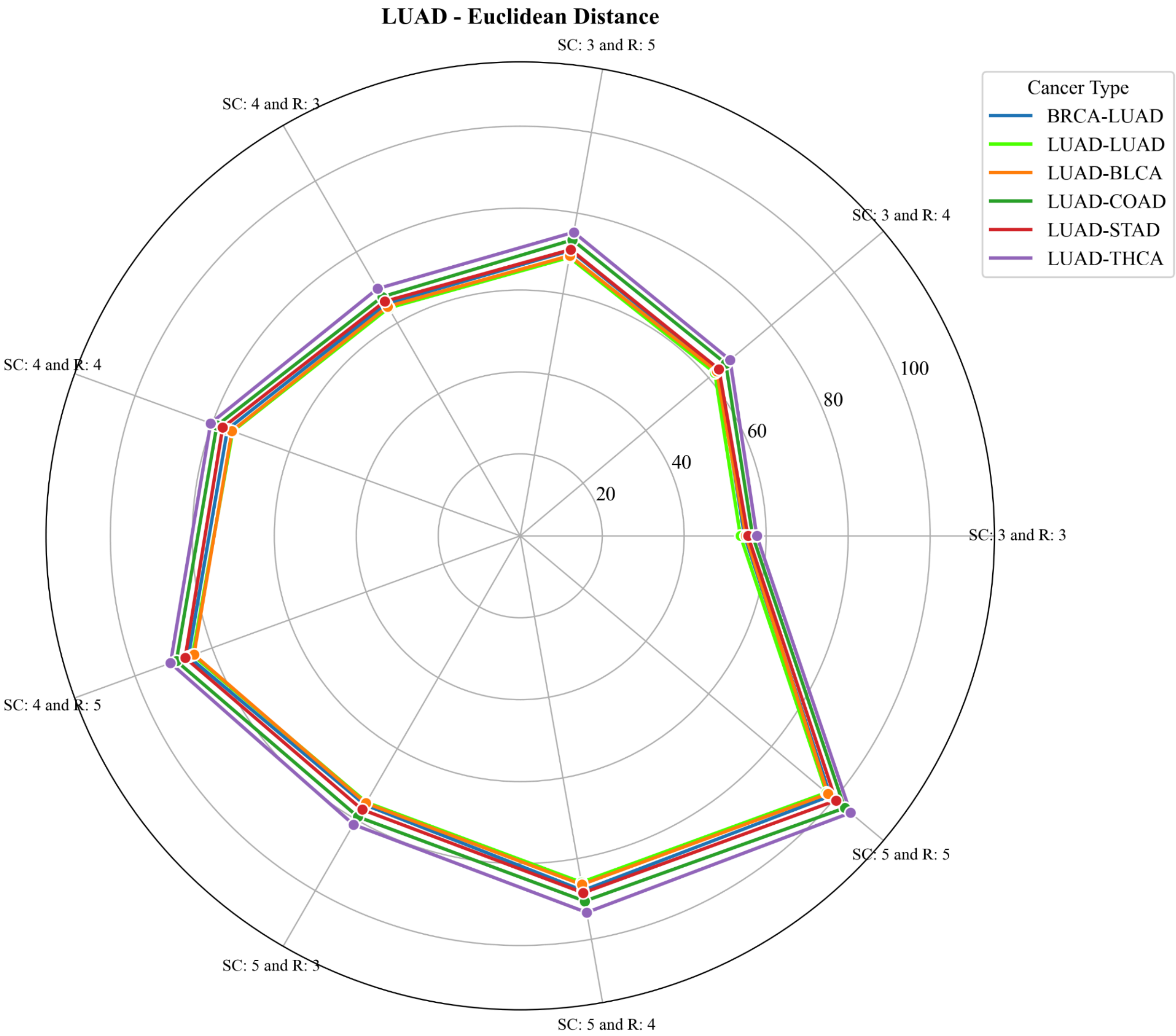
LUAD - Euclidean Distance
SC: 3 and R: 5
SC: 4 and R: 3
SC: 3 and R: 4
SC: 4 and R: 4
SC: 3 and R: 3
SC: 4 and R: 5
SC: 5 and R: 5
SC: 5 and R: 3
SC: 5 and R: 4
20
40
60
80
100
Cancer Type
BRCA-LUAD
LUAD-LUAD
LUAD-BLCA
LUAD-COAD
LUAD-STAD
LUAD-THCA

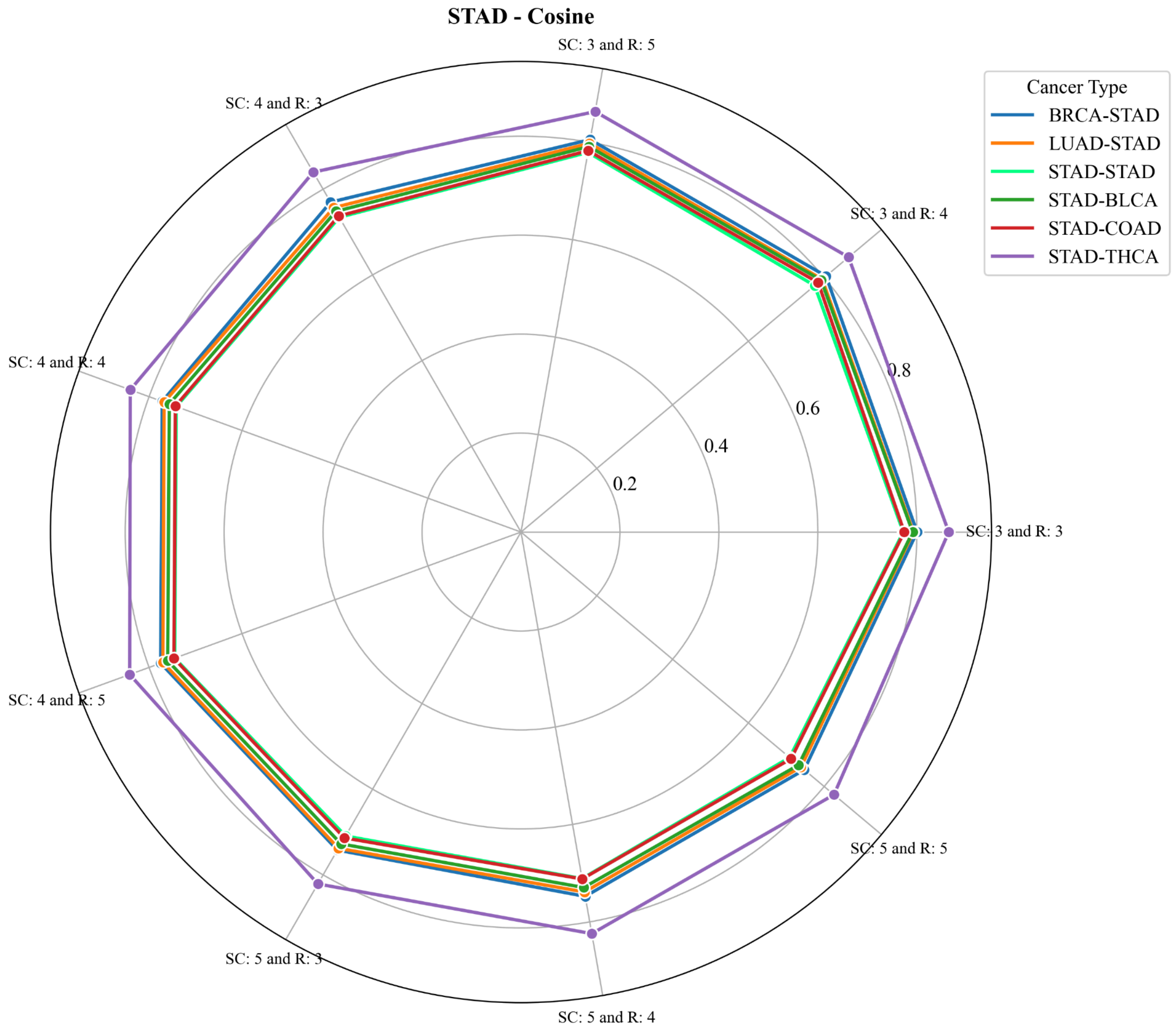
STAD - Cosine
SC: 3 and R: 5
SC: 3 and R: 4
SC: 3 and R: 3
SC: 5 and R: 5
SC: 5 and R: 4
SC: 5 and R: 3
SC: 4 and R: 5
SC: 4 and R: 4
SC: 4 and R: 3
0.2
0.4
0.6
0.8
Cancer Type
BRCA-STAD
LUAD-STAD
STAD-STAD
STAD-BLCA
STAD-COAD
STAD-THCA

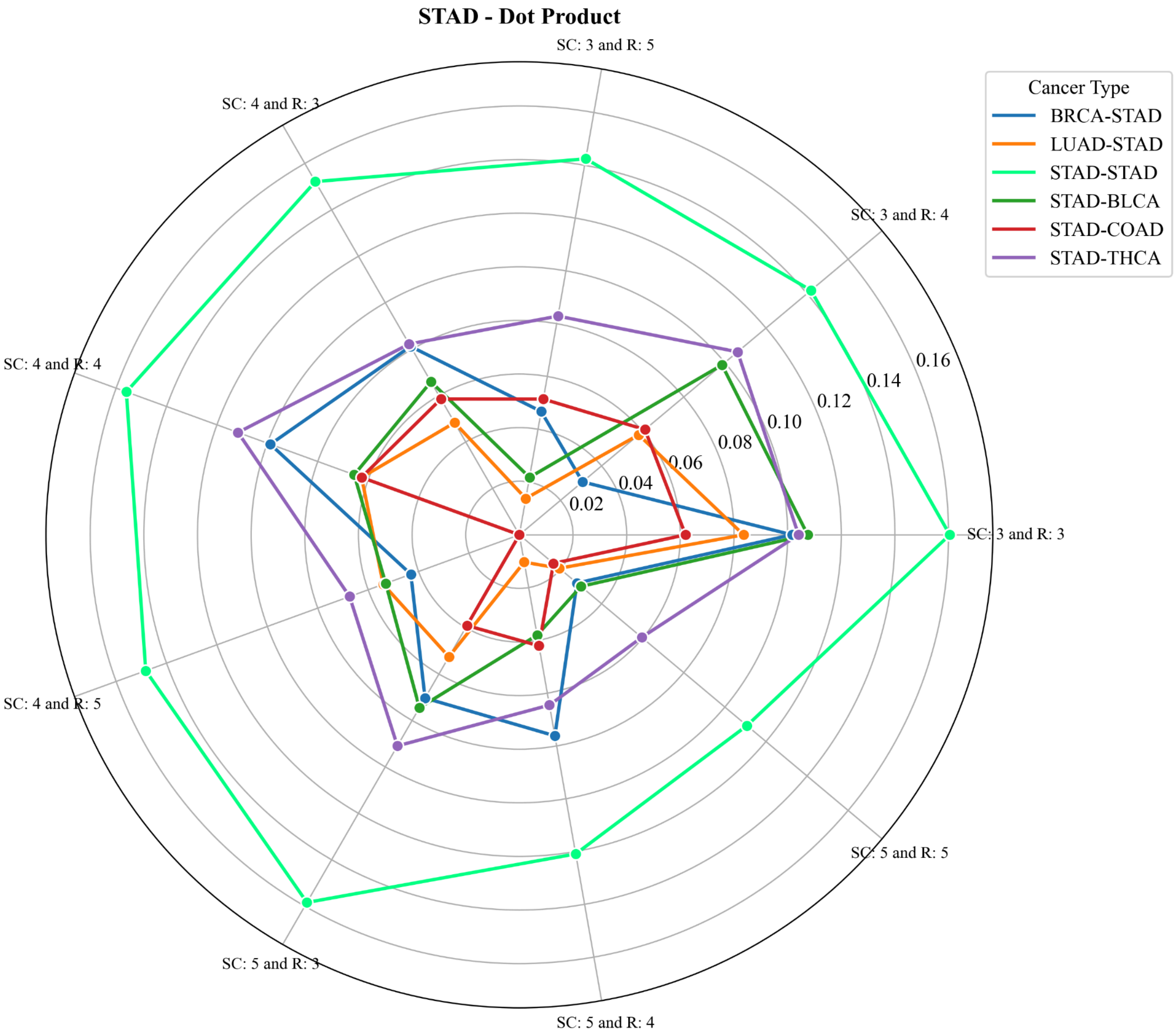
STAD - Dot Product
SC: 3 and R: 5
SC: 3 and R: 4
SC: 3 and R: 3
SC: 5 and R: 5
SC: 5 and R: 4
SC: 5 and R: 3
SC: 4 and R: 5
SC: 4 and R: 4
SC: 4 and R: 3
0.02
0.04
0.06
0.08
0.10
0.12
0.14
0.16
Cancer Type
BRCA-STAD
LUAD-STAD
STAD-STAD
STAD-BLCA
STAD-COAD
STAD-THCA

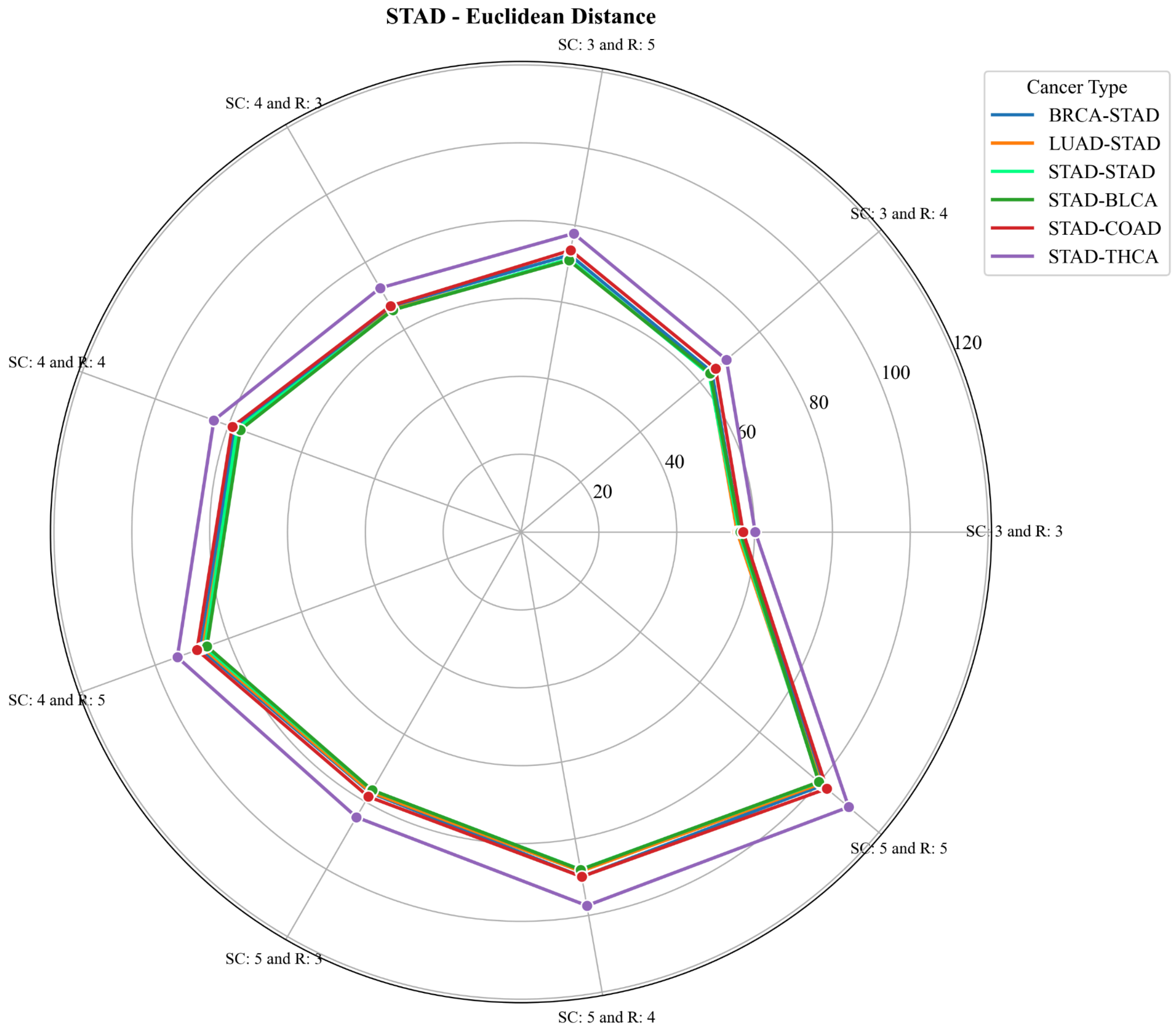

STAD - Euclidean Distance
SC: 3 and R: 5
SC: 4 and R: 3
SC: 3 and R: 4
SC: 4 and R: 4
SC: 3 and R: 3
SC: 4 and R: 5
SC: 5 and R: 5
SC: 5 and R: 3
SC: 5 and R: 4
20
40
60
80
100
120
Cancer Type
BRCA-STAD
LUAD-STAD
STAD-STAD
STAD-BLCA
STAD-COAD
STAD-THCA

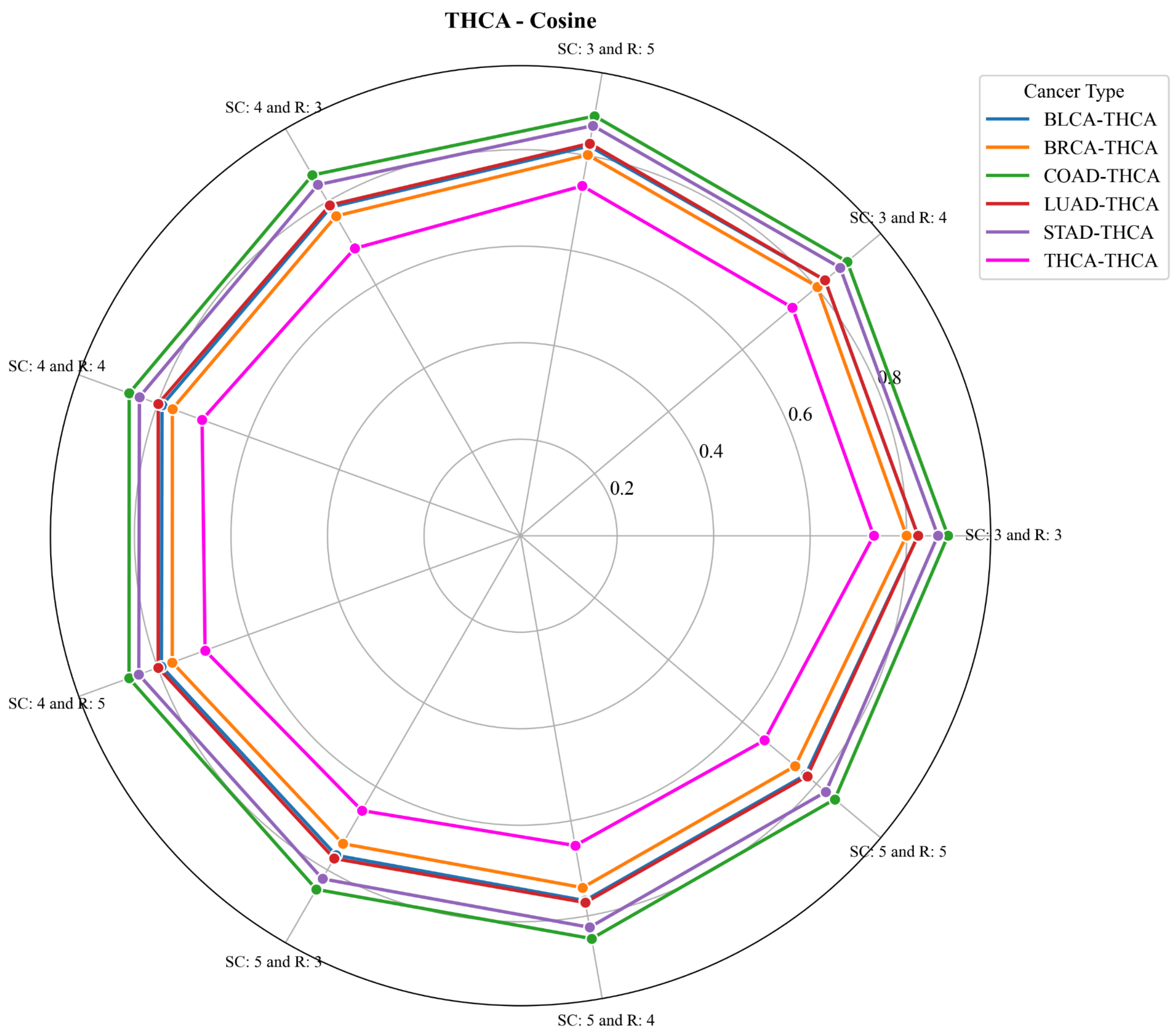
THCA - Cosine
SC: 3 and R: 5
SC: 3 and R: 4
SC: 3 and R: 3
SC: 5 and R: 5
SC: 5 and R: 4
SC: 5 and R: 3
SC: 4 and R: 5
SC: 4 and R: 4
SC: 4 and R: 3
0.2
0.4
0.6
0.8
Cancer Type
BLCA-THCA
BRCA-THCA
COAD-THCA
LUAD-THCA
STAD-THCA
THCA-THCA

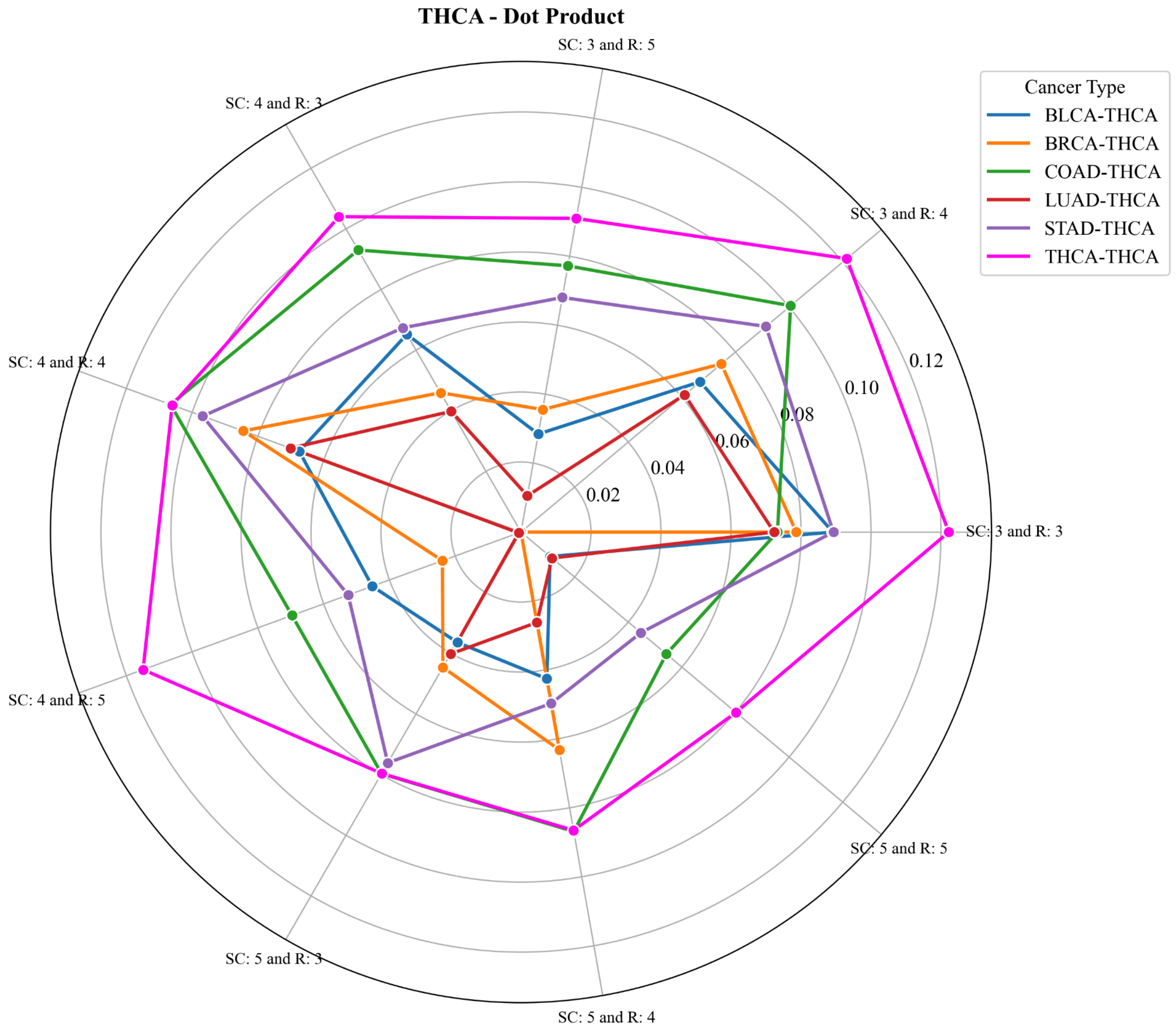

THCA - Dot Product
SC: 3 and R: 5
SC: 3 and R: 4
SC: 3 and R: 3
SC: 5 and R: 5
SC: 5 and R: 4
SC: 5 and R: 3
SC: 4 and R: 5
SC: 4 and R: 4
SC: 4 and R: 3
0.02
0.04
0.06
0.08
0.10
0.12
Cancer Type
BLCA-THCA
BRCA-THCA
COAD-THCA
LUAD-THCA
STAD-THCA
THCA-THCA

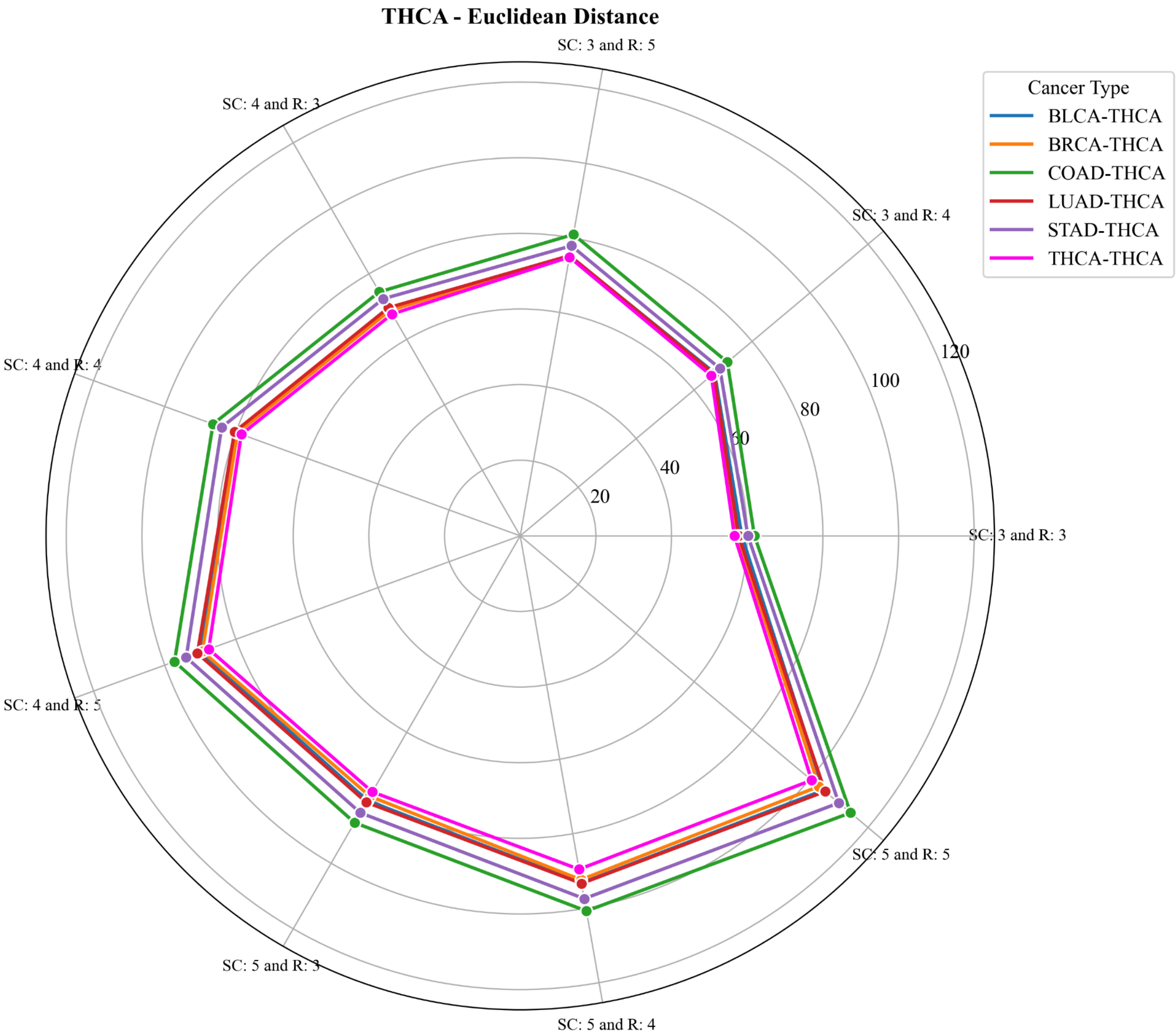


**Supplemental Figure 4**: Pairwise distance calculations between cancer types using Euclidean distance, cosine similarity, and dot product metrics demonstrate clear separation between intra- and inter-cancer distances. Cosine similarity and Euclidean distance are normalized between 0 and 1, whereas the dot product is only bounded at 0 with no defined upper limit.

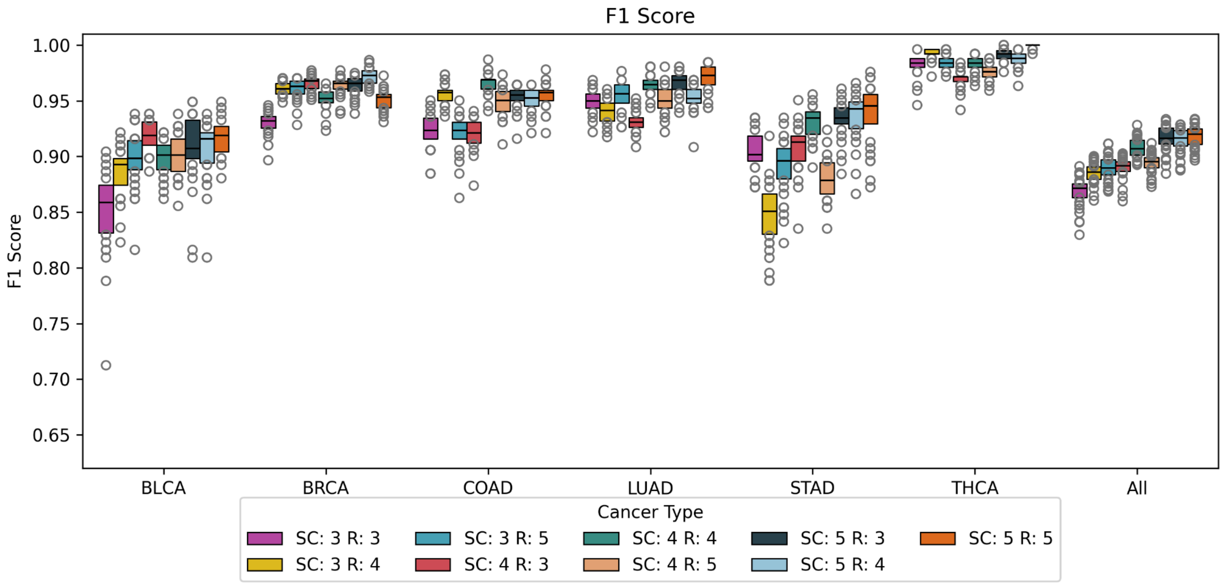


**Supplemental Figure 5**: F1 scores depicting classification performance across six different cancer types, evaluated over varying sample counts (SC) and repetitions (R), indicate that increasing either sample count or repetition enhances classification performance across all cancer types. The overall average F1 score across all cancers ranges between 0.90 and 0.95, demonstrating robust and consistent performance.

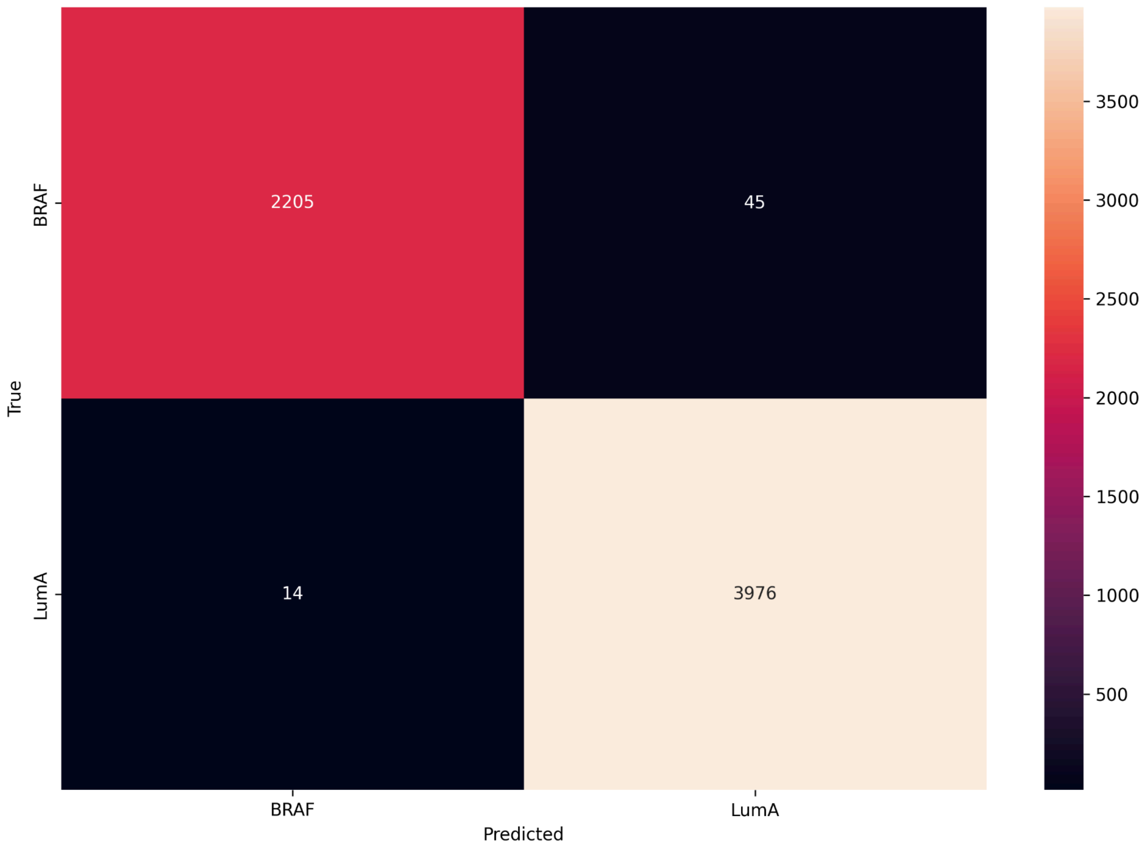


**Supplemental Figure 6:** Confusion matrix illustrating true versus predicted classifications for Luminal A (LumA) and BRAF cancer subtypes. Misclassifications are minimal, with a higher occurrence of BRAF samples being incorrectly classified as LumA.

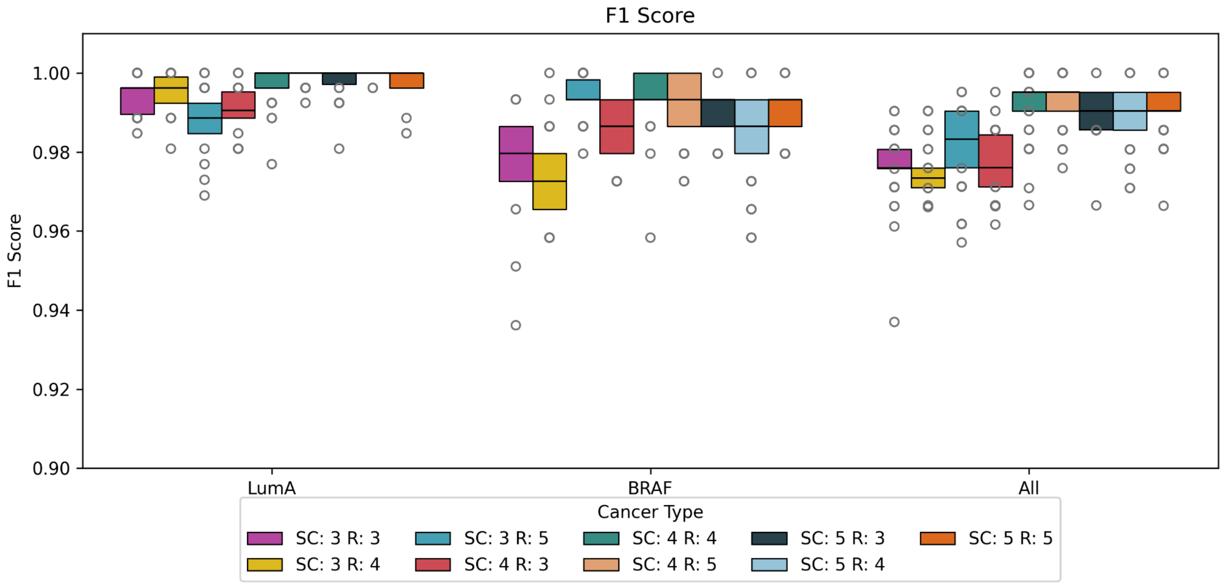

**Supplemental Figure 7:** F1 scores for subtype classification of Luminal A (LumA) and BRAF compared with overall F1 performance across varying sample counts (SC) and repeats (R), ranging from 3 to 5. Performance remains consistently high, with F1 scores ≥ 0.9 across all conditions.

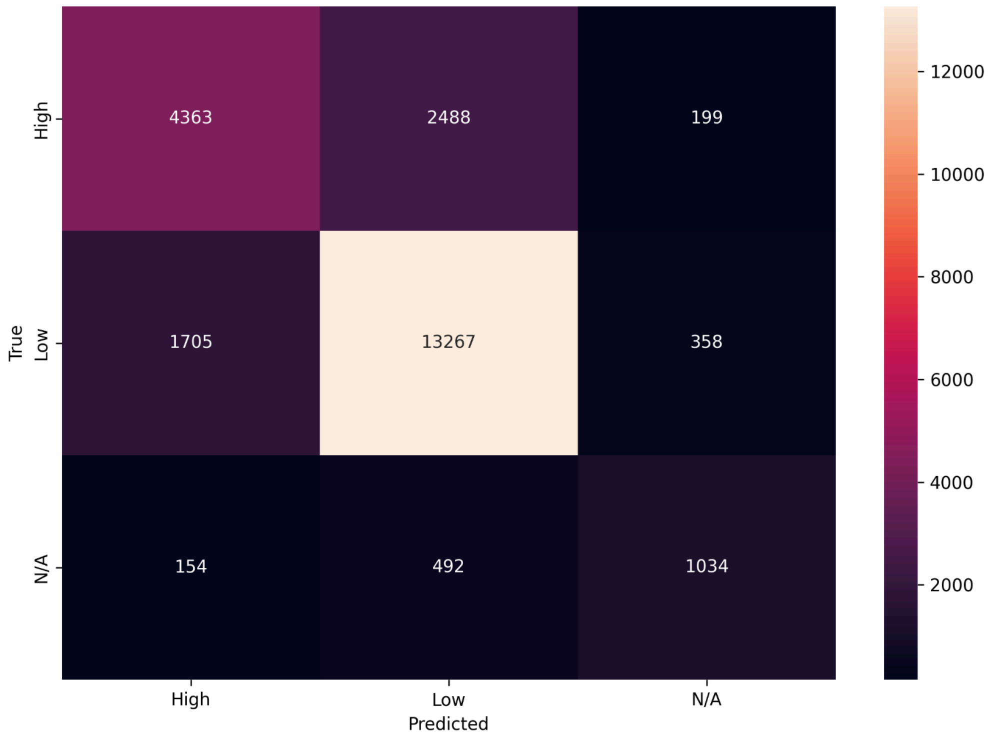


**Supplemental Figure 8**: Confusion matrix depicting true versus predicted classifications of Low, High, and N/A tumor mutational burden (TMB). Elevated misclassifications are observed for the N/A class, predominantly predicted as Low TMB. Additionally, minor misclassifications are noted for the N/A class when predicted as High TMB.

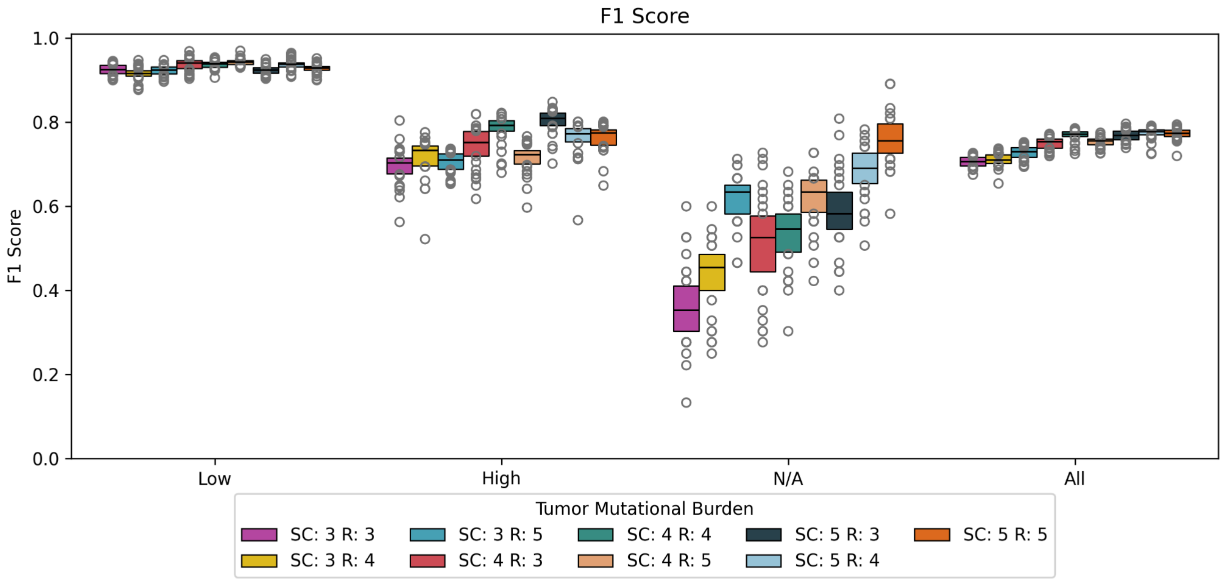


**Supplemental Figure 9:** F1 scores for Low, High, and N/A tumor mutational burden (TMB) classes, along with an overall average across all classes, are evaluated over varying sample counts (SC) and repetitions (R). The Low TMB class consistently achieves high performance with F1 scores ≥ 0.9, while the High TMB class maintains moderate performance ranging between 0.7 and 0.8. Notably, the N/A class exhibits poor performance with F1 scores < 0.4 when using a sample count of 3 and repetition of 3. However, increasing the sample count and repetition progressively improves N/A class performance, reaching an F1 score of approximately 0.8.
